# YhcB (DUF1043), a novel cell division protein conserved across gamma-proteobacteria

**DOI:** 10.1101/2020.12.31.425005

**Authors:** Jitender Mehla, George Liechti, Randy M. Morgenstein, J. Harry Caufield, Ali Hosseinnia, Alla Gagarinova, Sadhna Phanse, Mary Brockett, Neha Sakhawalkar, Mohan Babu, Rong Xiao, Gaetano T. Montelione, Sergey Vorobiev, Tanneke den Blaauwen, John F. Hunt, Peter Uetz

**Author notes:** Department of Chemistry and Biochemistry, University of Oklahoma, 101 Stephenson Parkway, Norman, OK 73019, United States. Correspondence: **Jitender Mehla**, **Peter Uetz**.

## Abstract

YhcB, an uncharacterized protein conserved across gamma-proteobacteria, is composed predominantly of a single Domain of Unknown Function (DUF 1043) with an N-terminal transmembrane α-helix. Here, we show that *E. coli* YhcB is a conditionally essential protein that interacts with the proteins of the cell divisome (e.g., FtsI, FtsQ) and elongasome (e.g., RodZ, RodA). We found 7 interactions of YhcB that are conserved in *Yersinia pestis* and/or *Vibrio cholerae*. Furthermore, we identified several point mutations that abolished interactions of YhcB with FtsI and RodZ. The *yhcB* knock-out strain does not grow at 45°C and is hypersensitive to cell-wall acting antibiotics even in stationary phase. The deletion of *yhcB* leads to filamentation, abnormal FtsZ ring formation, and aberrant septa development. The 2.8 Å crystal structure for the cytosolic domain from *Haemophilus ducreyi* YhcB shows a unique tetrameric α-helical coiled-coil structure that combines parallel and anti-parallel coiled-coil intersubunit interactions. This structure is likely to organize interprotein oligomeric interactions on the inner surface of the cytoplasmic membrane, possibly involved in regulation of cell division and/or envelope biogenesis/integrity in proteobacteria. In summary, YhcB is a conserved and conditionally essential protein that is predicted to play a role in cell division and consequently or in addition affects envelope biogenesis.

**Importance:** Only 0.8 % of the protein annotations in the UniProt are based on experimental evidence and thus, functional characterization of unknown proteins remains a rate-limiting step in molecular biology. Herein, the functional properties of YhcB (DUF1043) were investigated using an integrated approach combining X-ray crystallography with genetics and molecular biology. YhcB is a conserved protein that appears to be needed for the transition from exponential to stationary growth and is involved in cell division and/or envelope biogenesis/integrity. This study will serve as a starting point for future studies on this protein family and on how cells transit from exponential to stationary survival.

## Introduction

The sequencing revolution has flooded databases with millions of uncharacterized protein sequences. Only 0.8 % of the ~180 million protein sequences in UniProtKB/TrEMBL (1) are experimentally annotated or are associated with transcript data (0.72%) (Uniprot, Feb 2, 2020). Around 25.51% of sequence annotations have been inferred by homology and another 73.69% of sequences have been annotated by prediction algorithms (1). The functions of most proteins in Uniprot (or Pfam) are either computationally predicted or unknown. Therefore, functional characterization of unknown proteins, remains a rate-limiting step in molecular biology (2–4). Initially, *E. coli* YhcB was thought to be a subunit of cytochrome bd (oxidase) but was later found to be dispensable for the assembly of cytochrome bd (5). Large-scale genomic and proteomic studies indicated that *yhcB* may be involved in biofilm formation (6), cell envelope integrity (7), cold sensitivity (8), and DNA damage-associated (cell survival, repair) processes (9–11). Furthermore, a synthetic lethal phenotype was observed in combination with a cell shape maintenance gene deletion, *rodZ* (12). The latter study suggested a role in cell division which was recently confirmed by Sung et al. 2020 (13) who also found cell division defects in *yhcB* deletion strains. However, the molecular mechanism of these phenotypes remained unknown. Here, we investigate the structure and function of *E. coli* YhcB and its role in cell division by screening for YhcB-mutant phenotypes and for interacting proteins. Most importantly, we investigated the molecular basis for this function by determination of the x-ray crystal structure of the cytoplasmic region of the YhcB ortholog from *Haemophilus ducreyi*.

## Results

### YhcB is conserved in proteobacteria

YhcB is conserved across most gamma-proteobacteria but absent in other bacterial genomes (**Fig. 1**). The *yhcB* gene is located upstream of two periplasmic outer membrane stress sensor proteases (*degQ* and *degS*) and downstream of a cell division gene *zapE* (*yhcM*), which is encoded on the opposite strand (**Fig. S1**; for details, see legends of **Fig. S1**).

**Figure 1.**
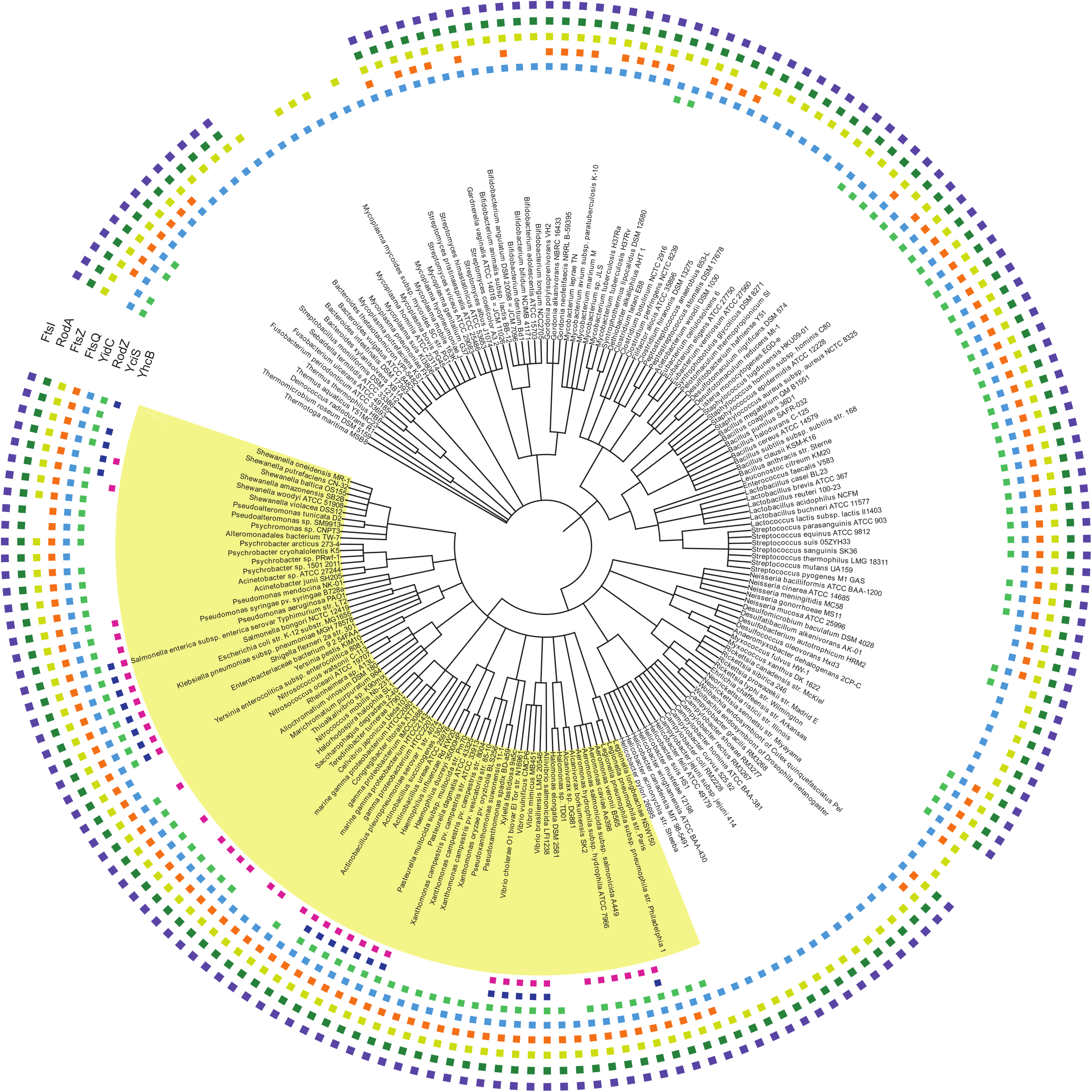
Phylogenomics of *yhcB* gene. Phylogenetic profile of YhcB and its interacting proteins. Proteobacteria are highlighted in red. *E. coli* is indicated by a white arrow.

### *yhcB* deletion results in multiple phenotypes

In order to understand the function and phenotypes of *yhcB*, we used a *yhcB* deletion strain to carry out extensive phenotyping. The Δ*yhcB* strain grows with a mass doubling time of 25 min, whereas the wild-type (WT) doubles every 22 min. Morphologically, cultures of *E. coli ΔyhcB* exhibited increased cell lengths but reduced diameters (**Fig. S2**). The Δ*yhcB* cells grow normal under exponential conditions but do not fully activate the growth arrest regulation towards stationary phase, which results in filaments. Stationary filamentous cells lacking *yhcB* exhibit no change in DNA concentration compared to WT cells (**Fig. 2A**), but DNA segregation is often disturbed. This is in contrast to exponential cells, where both DNA concentration and segregation appear to be unchanged in Δ*yhcB* cells compared to the WT strain (**Fig. 2B**). Additionally, Δ*yhcB* cells showed several other phenotypes, including temperature sensitivity (**Fig. 3A-B**; **Fig. S3a-d**). Given that we found cell division defects (e.g. filamentation,) and susceptibility of Δ*yhcB* strain to several PG-targeting antibiotics (**Fig. S4a),** (14) we tested the effect of two cell-wall targeting antibiotics (A22 and Mecillinam) on Δ*yhcB* cells. Proteins of the cell elongasome such as MreB and PBP2 are direct targets of the cell-wall antibiotics A22 and Mecillinam, respectively, and our experiments confirmed that the Δ*yhcB* strain was hypersensitive to both antibiotics (**Fig. 3C**).

**Figure 2.**
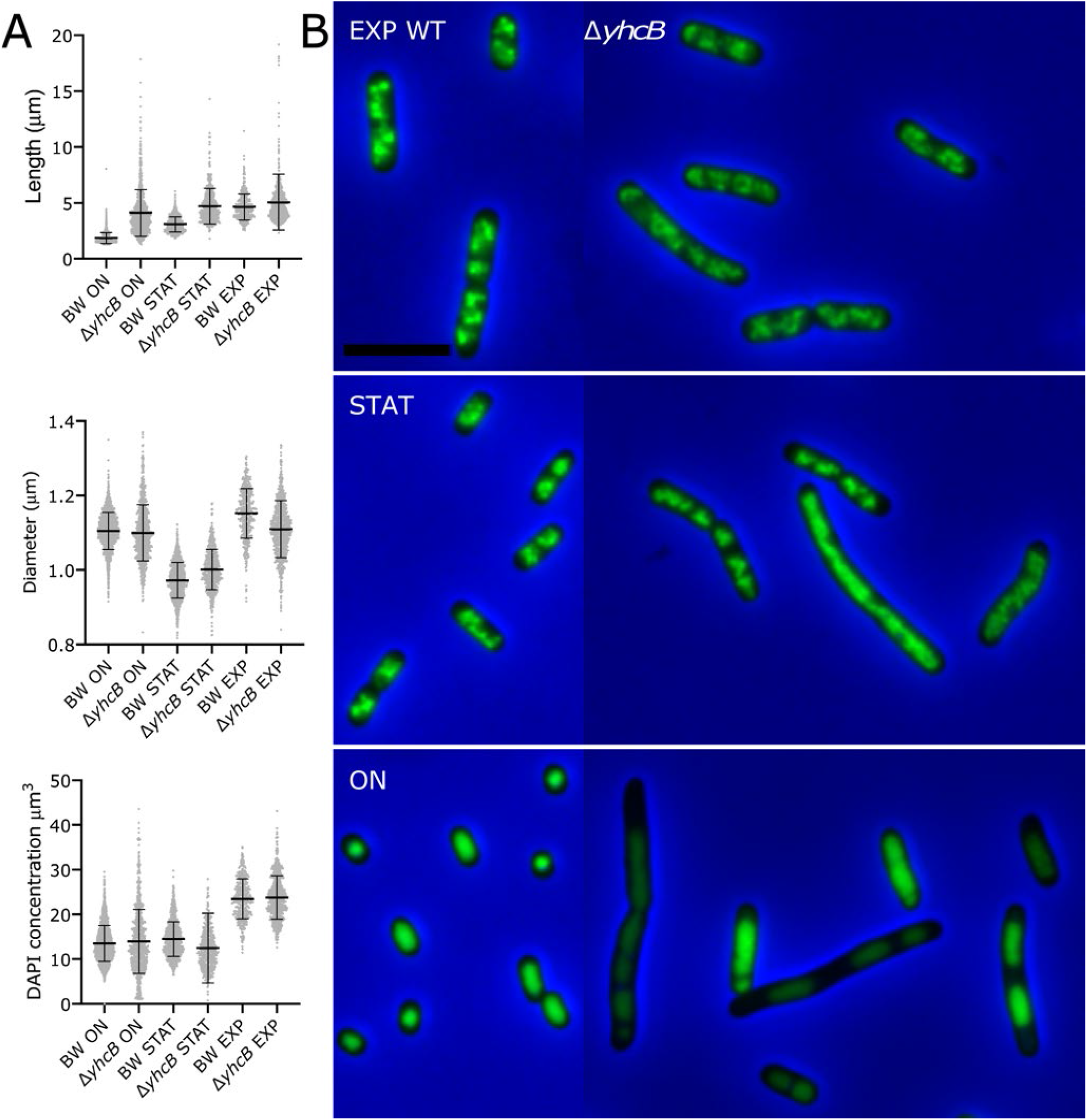
Δ*yhcB* lacks proper stationary state growth regulation. The Δ*yhcB* and its parental strain BW25113 (WT) were grown in LB at 37°C for 24 h (ON), then dilute 1:1000 and grown to an OD_650_ nm of 0.3 (EXP) or to and OD_650_ nm of 1.2 (STAT), fixed and the nucleoids were stained with DAPI. **(A).** length and, diameter and DAPI fluorescence of each culture with the mean and standard deviation indicated. BW EXP, STAT and ON number of analyzed cells were 434, 1225 and 2133 respectively, Δ*yhcB* EXP, STAT, and ON number of analyzed cells were 555, 776, and 811, respectively. **(B).** Representative images form all 6 cultures with BW25113 on the left and Δ*yhcB* on the right. The images are merged phase contrast (gray) and DAPI (green) images with a blue background for optimal contrast. Brightness and contrast are the same for all images. The scale bar equals 5 μm.

**Figure 3.**
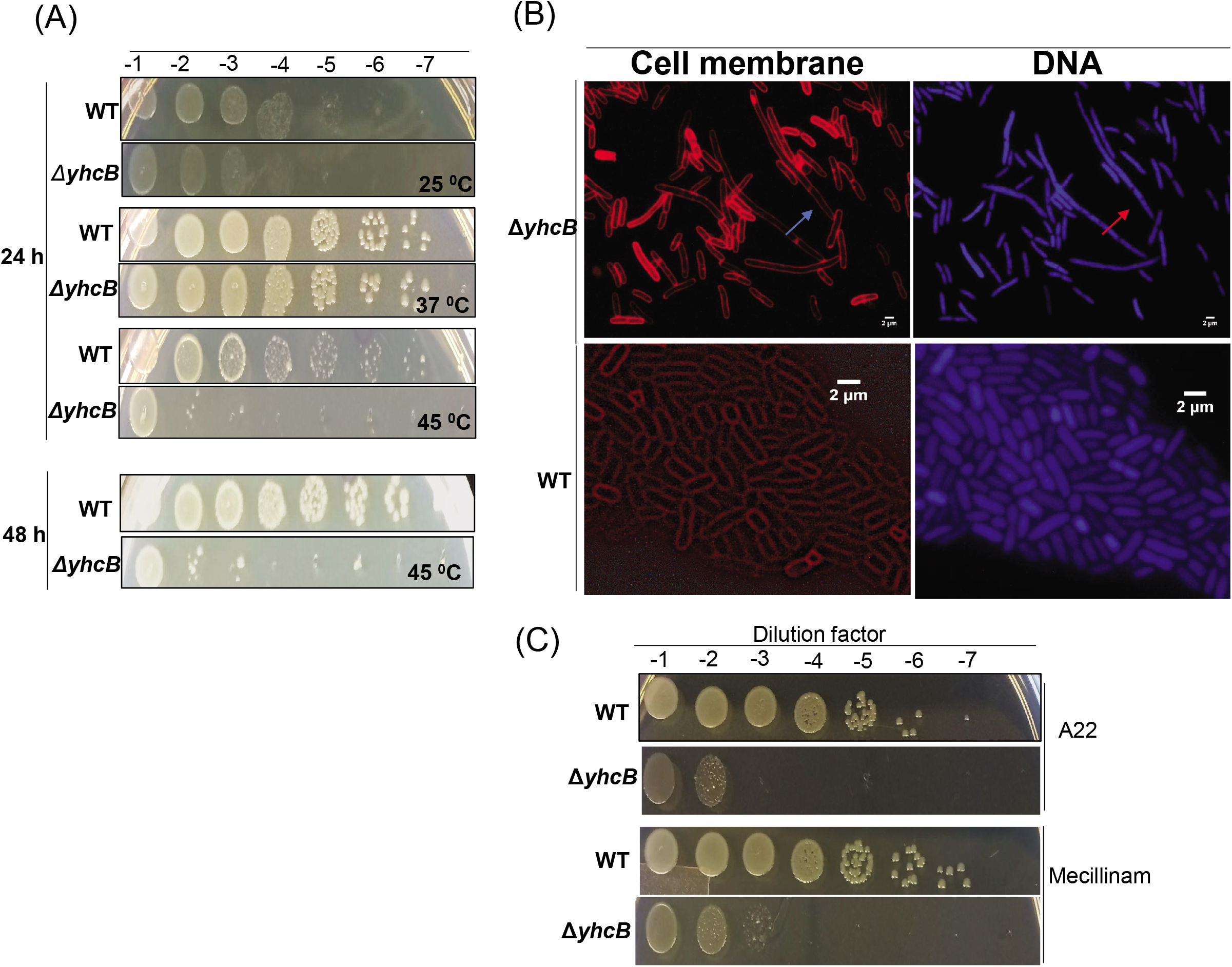
Δ*yhcB* mutant result in filamentation and susceptibility to antibiotics. **(A)** Temperature sensitivity of Δ*yhcB* cells. Δ*yhcB* cells are sensitive to high (45°C) temperature. (**B**) Micrographs of Δ*yhcB* cells in LB. The Δ*yhcB* cells with and without clear formation of septa were observed. (**C**) The Δ*yhcB* cells showed hypersensitivity of Δ*yhcB* cells to cell-wall acting antibiotics. Top: A22, bottom: Mecillinam.

We have not attempted to complement the aforementioned deletions by overexpression constructs, but such experiments have been described by Sung et al. 2020, showing that all phenotypes of their *yhcB* mutants were completely or significantly restored by YhcB expression (see Discussion for details).

### Stationary phase cultures of Δ*yhcB* strain exhibit susceptibility to cell-wall targeting antibiotics

Most of the inhibitors/antibiotics that target cell-envelope biogenesis, especially β-lactams, need actively growing cells to attain their maximum antibacterial activity. Given that a *yhcB* mutant strain exhibits hypersensitivity to antibiotics that target the bacterial cell wall (**Fig. 3C**), we tested Δ*yhcB* cells in early log phase, overnight and after 2 days in stationary phase. We counted a lower number of survivor cells in Δ*yhcB* strain compared to WT strain upon A22 treatment (**Fig. 4A**). No viable (persister) cell was observed after exposure of exponentially growing cells to Mecillinam (**Fig. 4A**). Two-day-old WT cells were least sensitive to cell-wall targeting antibiotics followed by overnight and exponential cells. However, we observed the opposite trend for the Δ*yhcB* cells in terms of their sensitivity towards cell-wall acting antibiotics. All mutant cells were found to be hypersensitive to the cell-wall antibiotics compared to overnight cells (**Fig. 4B)**. We also observed that WT cells adapted to antibiotic stress after 2 hours whereas Δ*yhcB* cells did not recover from the antibiotic shock even after 6 hours (**Fig. 4B**). A22 and Mecillinam inhibited growth of Δ*yhcB* mutants ~ 3-fold more than WT cells after 6 h (**Fig. 4B**). The hypersensitivity of 2 days old stationary cells indicates either an active PG synthesis machinery in stationary phase cells or defective cell envelope (**Fig. S4-b**). The latter was also supported by the β-galactosidase assay that reports envelope leakiness (**Fig. S3-c**).

**Figure 4.**
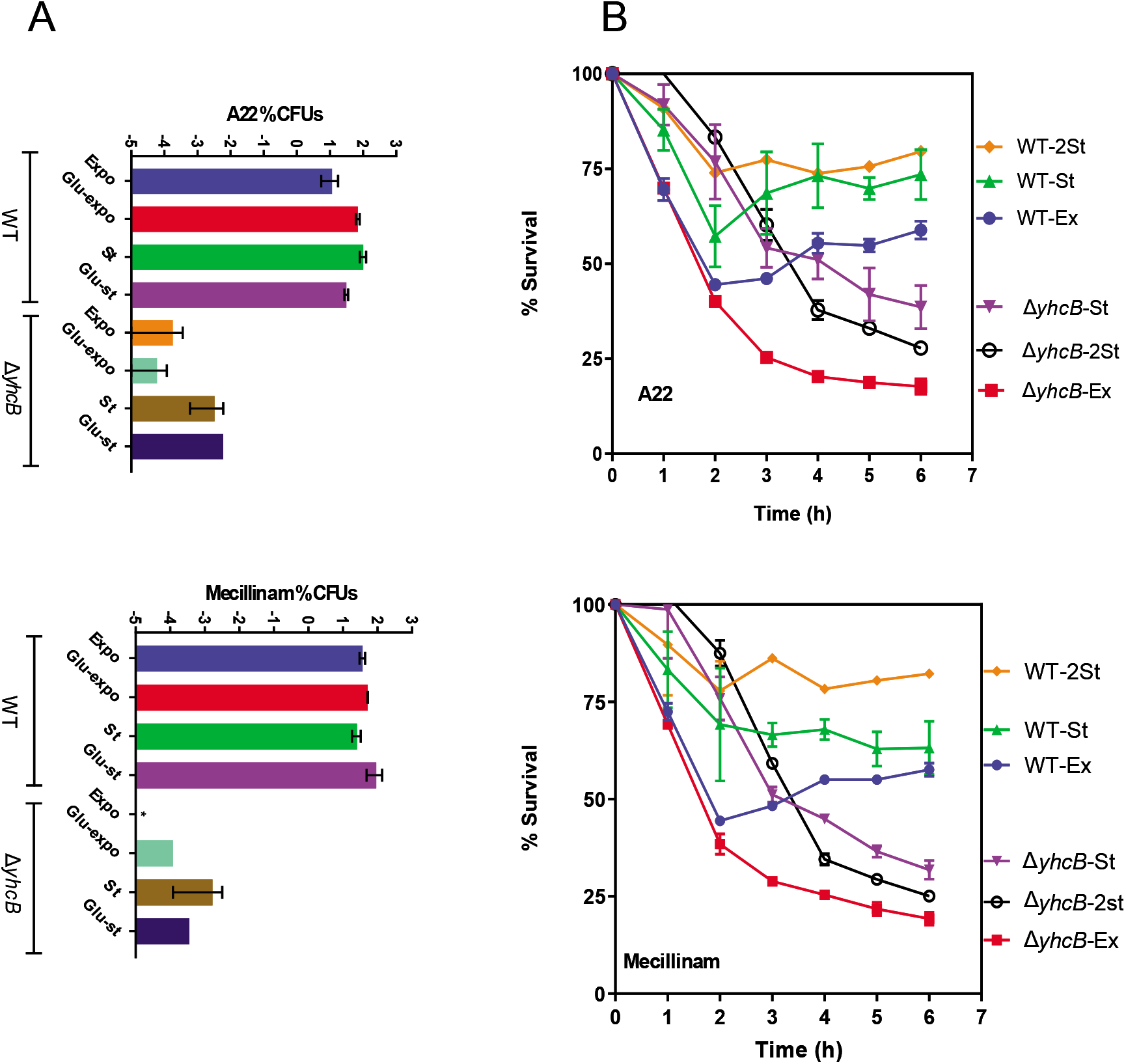
Hypersensitivity of Δ*yhcB* cells. **(A)** Fraction of surviving cells expressed as log% CFUs (**left**) and **(B)** % survival (**right**) of exponentially growing (“expo” or “ex”) and stationary phase cells (“st”) in LB media (2St = 2 days old stationary cells). Top: A22, bottom: Mecillinam.

### *yhcB* gene deletion leads to abnormal FtsZ ring and septum formation

The aforementioned Δ*yhcB* phenotypes indicate defective cell division in Δ*yhcB* mutant cells. Therefore, in order to visualize the cell membrane and clearly discern septum formation we stained the cells with SynaptoRed™C2 / FM4-64. No septum formation was observed in the majority of filamented cells (**Fig. 3B**). To determine if YhcB is necessary for successful formation of the bacterial divisome, we monitored FtsZ-ring formation in Δ*yhcB* cells. Immunolabelling with FtsZ-specific antibodies and secondary antibodies conjugated to a fluorophore in Δ*yhcB* cells showed that the Z-ring was not assembled properly/stably (**Fig. 5A-C**) despite sufficient concentration of FtsZ in Δ*yhcB* cells (**Fig. 5B**) and cells in all states potentially failed to form a Z ring. Notably, the Δ*yhcB* cells have more than twice the amount of FtsZ compared to the WT strain at the beginning of the stationary phase but that still did not rescue the phenotype. Furthermore, the FtsZ-ring formation appeared abnormal in the Δ*yhcB* strain with mis-localization of FtsZ (**Fig. 5D-E**, **Table 1)**.

**Figure 5.**
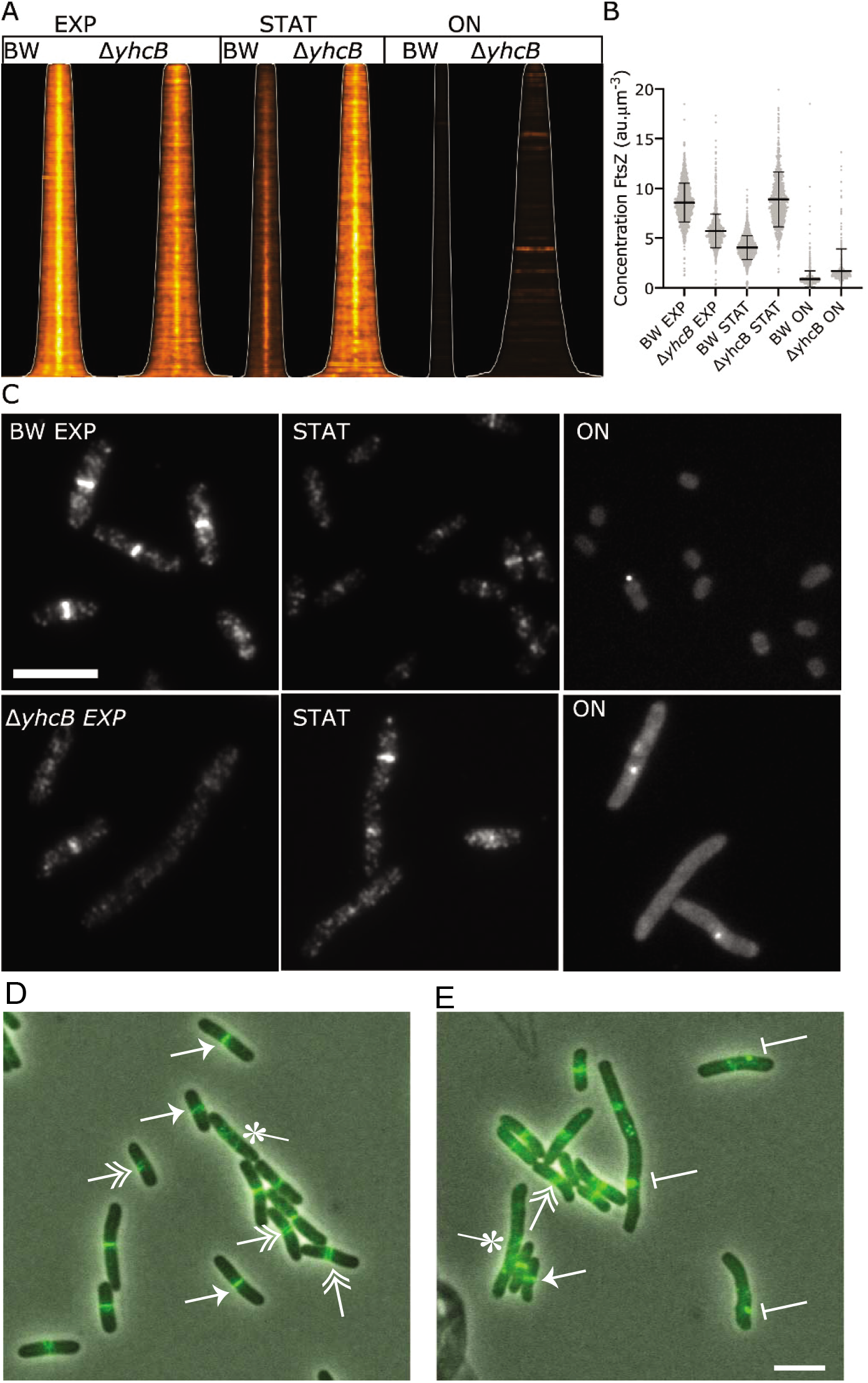
Δ*yhcB* cells display an increase in abnormal FtsZ localization. The Z-ring in Δ*yhcB* cells is not assembled properly as visualized by immunolabelling of FtsZ. (**A)** Map of FtsZ fluorescence profiles sorted according to cell length. The white line indicates where the cell poles are. Brightness and contrast are the same for all profiles. **(B)** The FtsZ concentration expressed in arbitrary units of all the cells of each culture with the mean and standard deviation indicated. The cells in EXP, STAT and ON phase (n=1735, 1606 and 1321, respectively) were analyzed for BW and Δ*yhcB* cells (n=1154, 964, and 721) respectively. **(C)** Representative fluorescence images from all 6 cultures with BW25113 (top) and Δ*yhcB* (bottom). The brightness and contrast of the images of the EXP and STAT cell is 0/13000 whereas it is 0/1300 for the images from the ON cells. The scale bar equals 5 μm. **(D)** WT cells expressing FtsZ-GFP^sw^. **(E)** Representative image of Δ*yhcB* cells expressing FtsZ-GFP^sw^. Different classes of FtsZ localization are indicated as follows: arrow-head-Z-ring, double arrowhead-helix, star-diffuse, bar= bright foci.

**Table 1.** FtsZ localization in WT or Δ*yhcB* cells. WT pattern contains cells that showed a central Z-ring or helix. Diffuse indicates that cells did not show any discernable pattern of FtsZ localization. Abnormal indicates cells with bright foci, multiple Z-rings, or off center Z-rings. Error is 90% confidence interval.

| Strain | Total Cells | WT pattern | Diffuse | Abnormal |
| --- | --- | --- | --- | --- |
| RM586 (WT) | 675 | 88% ± 4.6 | 10% ± 0.41 | 2% ± 0.10 |
| RM588 ($\Delta yhcB$) | 793 | 72.4 % ± 3.4 | 15.4% ± 0.60 | 12.2% ± 0.50 |

### Peptidoglycan (PG)-labelling showed incomplete septa and absence of septal PG formation in Δ*yhcB* filaments

The Δ*yhcB* strain showed impaired FtsZ ring formation, defective cell division, and hypersensitivity to antibiotics that target the cell wall (e.g. PG synthesis). Therefore, to locate intracellular sites of YhcB activity, we sought to monitor peptidoglycan (PG) synthesis in Δ*yhcB* cells. PG-labeling in a Δ*yhcB* strain was probed using a non-toxic, fluorescent D-amino acid analog of D-alanine (NADA), (15), which incorporates into the stem peptide of previously synthesized PG in living bacteria (**Fig. 6A**). In addition, we used another modified, D-amino acid dipeptide, EDA-DA (15–17) that incorporates specifically into the stem peptide of newly synthesized PG in the bacterial cytoplasm (**Fig. 6B**). Utilizing both probes, we are able to investigate whether the processes of PG synthesis and turnover were significantly affected in our Δ*yhcB* strain, as well as observed defects in septum formation.

**Figure 6.**
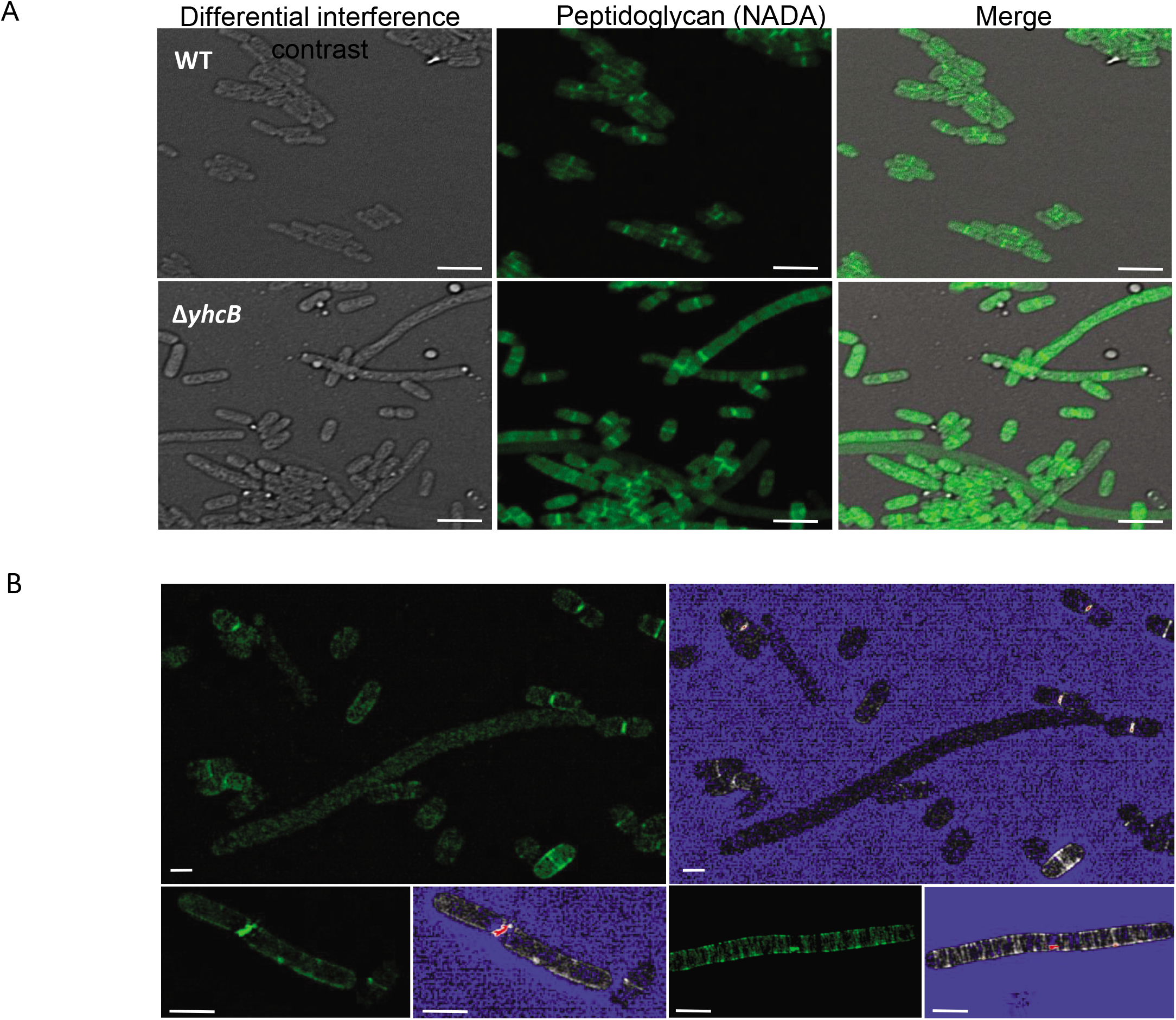
YhcB affects peptidoglycan localization and septum formation. (**A)** Wild type *E. coli* and Δ*yhcB* mutant cells subjected to a short (45 second) labeling pulse with the fluorescent D-alanine analog, NADA. Septa are observable within the smaller, more ‘wild-type’-looking Δ*yhcB* cells, while few visible septa are visible in elongated cells. (**B)** Structured Illumination microscopy (SIM) of the Δ*yhcB* mutant strain labeled with the D-alanine dipeptide analog, EDA-DA. Long, filamentous morphotypes are shown that appear to lack probe incorporation (indicative of an absence of newly forming septa, top panel) or exhibit abnormal, ‘punctate’ labeling, similar to FtsZ labeling shown in **Fig. A**, (bottom panels). Green panels show images as they appear in the FITC channel and blue panels show corresponding fluorescence intensity maps that range pixel intensities between 0 (blue) to 255. Scale bars; ~1 μm.

Not surprisingly, we observed far fewer labeled septa in the elongated forms of Δ*yhcB*. Similar to our previous observations, we also noticed a population of WT-like cells (in terms of length and presence of labeled division septa). For the filamented forms, we observed what appeared to be septal labeling using both NADA and EDA-DA probes, however, septum formation often appeared either aberrant or incomplete (**Fig. 6B).** We did observe PG labeling around the cell periphery in some elongated cells, indicating that new PG synthesis by the elongasome appears to occur in these cells for some period of time. In conclusion, peptidoglycan synthesis seemed to function apart from septum synthesis in filaments with diffuse Z-rings.

### YhcB genetically interacts with proteins of the cell division apparatus

Given that *yhcB* is responsible for several phenotypes, we investigated the epistatic connections of *yhcB* with other bacterial genes (i.e. if phenotypes of one mutation are modified by mutations in other genes). For that purpose, we used data from our previous envelope integrity study of *Escherichia coli* screened under both auxotrophic (rich medium) and prototrophic (minimal medium) conditions. Strikingly, at a high stringent filtering of the genetic interaction score (|E-score|≥10; *P* ≤ 0.05; **Table 2**; **Fig. 7A**), except *ftsE and rodZ*, we found 28 condition-dependent synthetic lethal interactions for gene pairs involved in cell division, cell shape, and cell wall biogenesis (or integrity), indicating that these genes are functionally related.

**Figure 7.**
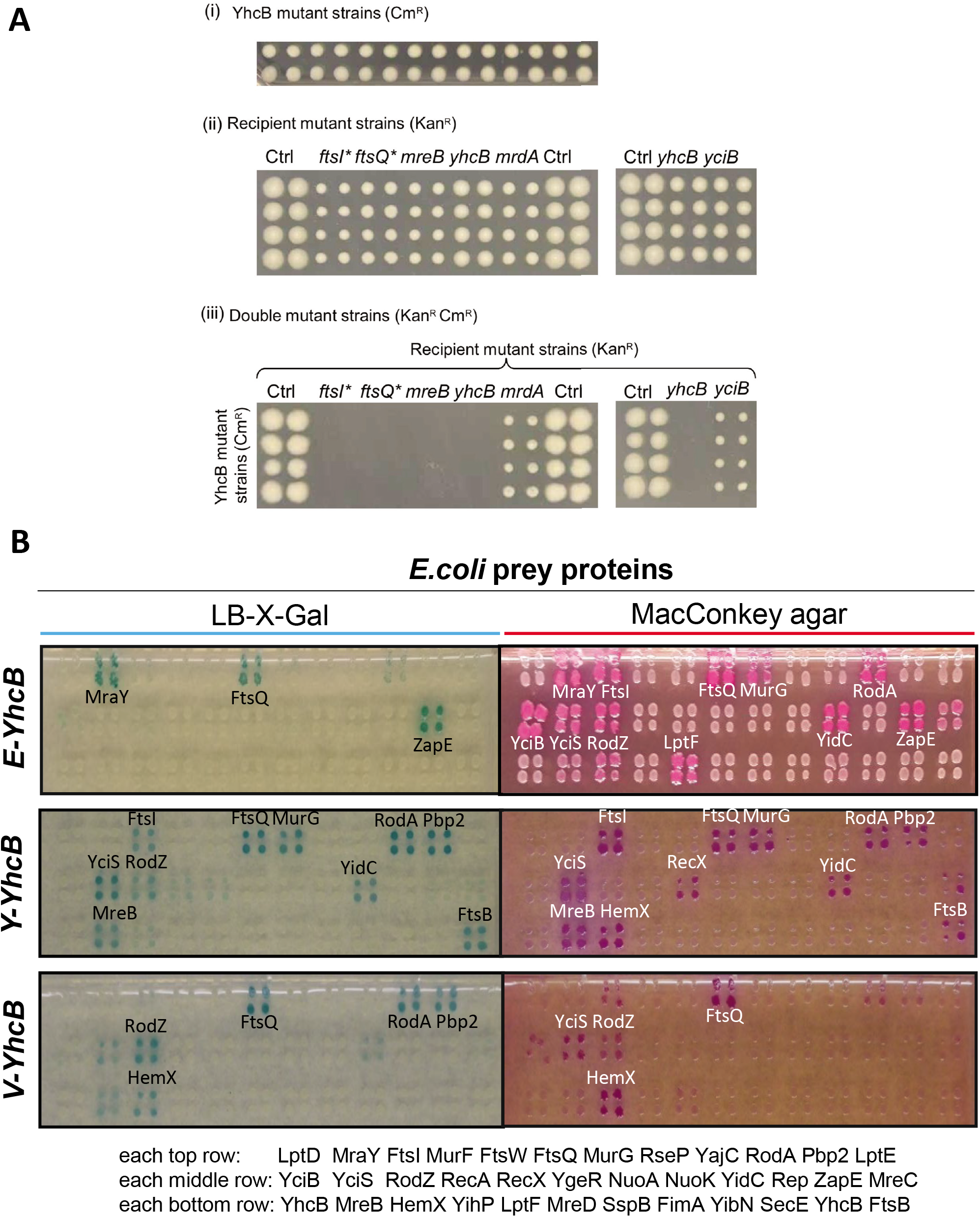
Interactions of *yhcB*. **(A)** Double mutants (iii) generated in rich medium by conjugating the Hfr Cavalli (HfrC) *yhCB* with chloramphenicol (Cm^R^) resistance (i) and the indicated F-recipient non-essential single gene deletion or essential hypomorphic (asterisk) mutant strains (ii) marked with kanamycin resistance (Kan^R^) marker. **(B)** A representative B2H screen of YhcB of *E. coli, Y. pestis and V. cholerae* against *E. coli* prey proteins. The colored colonies showed positive interactions. The percentage shows the identity between *E.coli* YhcB vs *Yersinia* and *Vibrio* YhcB. See text and methods for details.

**Table 2.** *yhcB* synthetic lethal genetic interaction pairs in cell division, cell shape, and cell wall biogenesis. E-RM = E-ScoreRM (rich media) and E-MM (minimal media) indicate synthetic lethal GIs, with “S” indicating strong synthetic lethal effects. See text and methods for details.

| Gene | E-RM | E-MM | Function |
| --- | --- | --- | --- |
| BcsB |  | S | Cell Shape, Glycan metabolism |
| CsrA |  | S | Cell Shape |
| DacA |  | S | Cell Wall Biogenesis |
| DacB |  | S | Cell Wall Biogenesis |
| DacC |  | S | Cell Wall Biogenesis |
| DdpC |  | S | Cell Wall Biogenesis |
| DdpF |  | S | Cell Wall Biogenesis |
| FtsA |  | S | Cell Division, Cell Shape |
| FtsE | S | S | Cell Division |
| FtsK |  | S | Cell Division |
| FtsZ |  | S | Cell Division, Cell Shape |
| GlmU |  | S | Cell Wall Biogenesis |
| ManY | S |  | Cell Wall Biogenesis |
| MepA |  | S | Cell Wall Biogenesis |
| MipA |  | S | Cell Wall Biogenesis |
| MraY |  | S | Cell Wall Biogenesis |
| OppC | S |  | Peptide transport |
| OppD |  | S | Peptide transport |
| PgpB |  | S | Cell Wall Biogenesis |
| Prc | S |  | Cell Division, Cell Wall Biogenesis |
| PtsH | S |  | Sugar transport |
| PtsI | S |  | Sugar transport |
| RodZ | S | S | Cell Shape, Cell Wall Biogenesis |
| RsmG |  | S | rRNA processing |
| Slt |  | S | Cell Division, Cell Wall Biogenesis |
| YehU |  | S | Cell Wall Biogenesis |
| YfeW | S |  | Cell Wall Biogenesis |
| YgeR |  | S | Cell Division |
| ZapB |  | S | Cell Division |
| ZipA |  | S | Cell Division, Cell Shape |

### YhcB co-purifies with cell division proteins

Next, we searched for YhcB interacting partners by expressing the protein with a C-terminal affinity from its native locus to maintain physiological protein level. YhcB was then affinity-purified (AP) from detergent solubilized cell extracts and analyzed by mass spectrometry (MS). In addition, we gathered proteins associated with YhcB in previous AP/MS and co-fractionation studies (18), as well as from quantitative proteomics (19) without epitope tagging. By combining these four sets of data, we were able to identify 49 high-confidence proteins that co-purified with YhcB and are involved in cell division / shape / biogenesis or maintaining membrane barrier function (**Table S1**).

### Binary protein-protein interactions of YhcB

Based on the interactions we found for YhcB from the above proteomic screens, as well as their relevance to *yhcB* phenotypes (e.g. RodZ), and results from other literature/database surveys, we chose 35 candidate proteins to test for direct interaction with YhcB (**Table S1**) using a bacterial two hybrid (B2H) system (20). We were able to verify a total of 10 interactions in *E. coli* (**Table 3**) that were detected in multiple assays and/or conserved across species. Six of those were confirmed by the aforementioned MS-based proteomics dataset **(Table S1)**, consistent with the validation rate typically observed for *E. coli* proteins using B2H assays (18, 21).

**Table 3.** Protein-protein interactions of YhcB in *E. coli, Y. pestis*, and *V. cholera*, based on Bacterial Two Hybrid screening (see Methods for details). Green boxes (Y) indicate positive interactions. The interaction with HemX (yellow) is conserved in all three species. GI = genetic interaction (see text for details). For cross-species interactions see **Table S2**.

| **Baits:** | **E-YhcB** | **Y-YhcB** | **V-YhcB** | **GI** | **MS detection (18)** |
| --- | --- | --- | --- | --- | --- |
| **Preys** | *E. coli* | *Yersinia* | *Vibrio* | *E. coli* | *E. coli* |
| degQ |  | Y |  |  |  |
| degS |  |  | Y |  |  |
| FtsA |  | Y |  | X |  |
| **FtsB** |  | Y | Y |  |  |
| FtsI | Y |  |  |  | Y |
| **FtsQ** | Y | Y |  |  | Y |
| FtsZ |  | Y |  | X | Y |
| **HemX** | Y | Y | Y |  |  |
| LptF |  | Y |  |  |  |
| MreB |  | Y |  |  |  |
| MurF |  |  | Y |  |  |
| MurG | Y |  |  |  |  |
| **RodA** | Y |  | Y |  |  |
| **RodZ** | Y | Y |  | X | Y |
| **YciB** | Y | Y |  |  |  |
| **YciS** | Y | Y |  |  | Y |
| **YidC** | Y | Y |  |  | Y |
| ZapB |  | Y |  | X |  |
| ZapE | Y |  |  |  |  |

In order to find biologically relevant and conserved interactions, we also tested the interactions found among *E. coli* proteins with their homologs from *Yersinia pestis* and *Vibrio cholerae* (**Fig. 7B**, see also **Fig. 1**). We detected 13 and 5 interactions of YhcB in *Yersinia pestis* and *Vibrio cholerae*, respectively (**Table 3**). Six *Yersinia* and two of the *Vibrio* interactions were also detected in *E. coli* (**Table 3**). Interactions that were detected in at least two species were considered to be conserved (and thus as more reliable) and we found 8 interactions in at least 2 species (**Table 3**). Only one interaction was detected in all three species, that of YhcB with HemX (**Table 3**).

We also tested cross-species interactions, that is, YhcB of *E. coli, Y. pestis* and *V. cholerae* were tested against test proteins of *E. coli, Y. pestis*, and *V. cholerae* for both intra-and inter-species interactions (**Table S2**). For instance, 4 YhcB interactions were found between *E. coli* YhcB and *V. cholerae* MurF, RodA, ZapE, and HemX, respectively, although YhcB shares only 45% sequence identity with its orthologs in both species. In addition, 8 PPIs were found between *E. coli* and *Y. pestis* which share 80% identity between their YhcB proteins (**Table 3**), and a few more across various combinations of the three bacteria (**Table S2**).

Importantly, YhcB interacts physically with proteins that comprise the cell elongasome (e.g. **RodZ, RodA**) and divisome (e.g. **FtsI, FtsQ**), complexes that are involved in cell-wall biogenesis and septum formation. Consistent with this observation, in addition to a *rodZ* mutant, we were able to confirm synthetic lethal or loss of fitness interactions between *yhcB* and genes involved in cell division (e.g. *ftsI, ftsQ*) cell-wall biosynthesis (*mrdA*), and cell shape maintenance (e.g. *mreB*) (**Fig. 7A-B**). These observations provide strong genetic and physical evidence that YhcB is involved in cell division and / or cell-wall biogenesis.

### Crystal structure of the YhcB cytoplasmic domain

In order to reveal the molecular basis of YhcB function, we determined its crystal structure. Screening of several proteobacterial orthologs for their purification and crystallization behavior led to us to focus on the structure determination of the cytoplasmic region of YhcB from the gamma proteobacterium *Haemophilus ducreyi*, an opportunistic genital pathogen. We expressed a truncated version of 132 amino acid protein in *E. coli* with a deletion of the predicted N-terminal transmembrane α-helix (residues 2-30) (22). A hexahistidine affinity tag was added to its native C-terminus for purification. We performed hydrodynamic analyses on the crystallization stock of this purified protein construct using size exclusion chromatography with multi-angle light scattering (SEC-MALS), which showed that it is primarily monomeric but forms small amounts of stable tetramer (2.5%) and hexadecamer (0.9%) in solution (**Fig. S5**).

This cytosolic region produced crystals that diffracted to ~3 Å resolution, but they could not be solved using anomalous diffraction from selenomethionine-labeled protein due to the absence of any internal methionine residues in the native protein sequence. We therefore introduced I51M and L72M mutations at two conserved hydrophobic sites that have methionine in some YhcB orthologs, which enabled us to solve and refine the structure at 2.8 Å resolution using single-wavelength anomalous diffraction from selenomethionine-labeled protein (**Table S3** and **Fig. 8**). Validation of the crystal structure is described in the Methods section.

**Figure 8.**
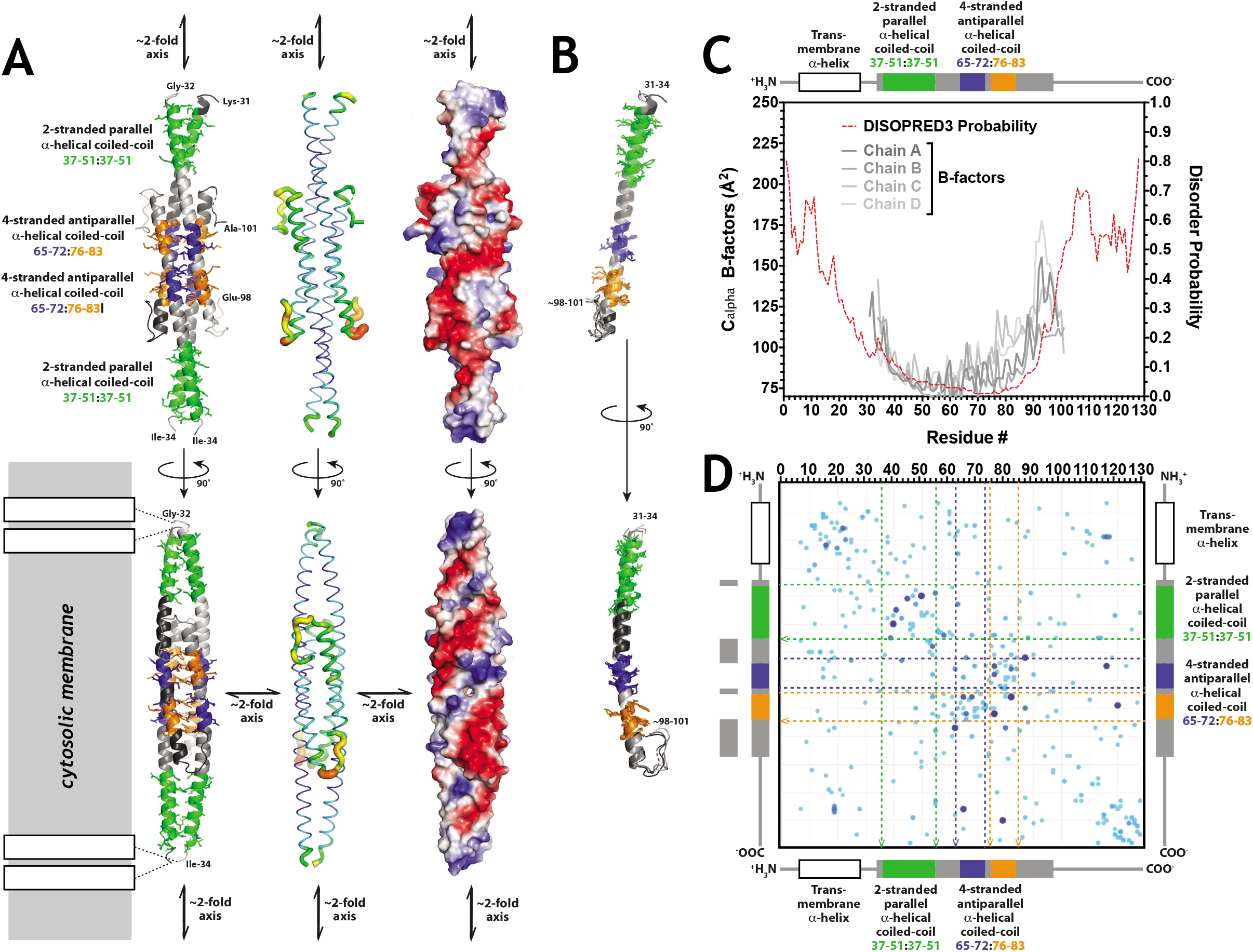
Crystal structure of the YhcB ortholog from *Haemophilus ducreyi*. **(A)** Ribbon diagrams (left), B-factor-encoded backbone traces (center), and surface electrostatic representations of two views related by a 90° rotation around the long axis of the coiled-coil homotetramer in the asymmetric unit of the crystal structure. The green and blue/orange colors in the ribbon diagrams show, respectively, the segments participating in parallel and antiparallel coiled-coil interactions in the tetramer. The rectangles with black borders at bottom left schematize the approximate geometry of the predicted N-terminal transmembrane □-helix deleted from the crystallized construct. The blue/narrow to red/wide gradient in the B-factor-encoded backbone traces span 74-174 Å^2^. The fully saturated blue/red colors on the molecular surfaces encode vacuum electrostatic potentials of +/- 93 kT calculated using the default parameters in PyMOL. **(B)** Ribbon diagrams showing least-square superposition of the four individual subunits in the asymmetric unit of the crystal structure, which are colored according to parallel *vs*. antiparallel coiled-coil interaction as in the leftmost images in panel A. **(C)** The backbone B-factors in the four subunits in the crystal structure (gray traces) plotted along with the probability of backbone disorder (red trace) calculated by the program DISOPRED3 (23) from the YhcB sequence profile. The 2° structure and parallel*/*antiparallel coiled-coil interactions observed in the crystal structure are schematized above the plot using the same color-coding as in the leftmost images in panel A. **(D)** Plot of pairwise evolutionary couplings (80) or probability of correlated evolutionary variations in the sequences of YhcB orthologs. The strength and statistical significance of each pairwise coupling is proportional to the diameter and darkness of the blue color of the circles, which represent *p*-values from 0.6-1.0 (scaled scores from 1.0-2.7) calculated using ~2.4 sequences per residue.

The crystal structure of the cytosolic region of *H. ducreyi* YhcB shows a coiled-coil tetramer (**Fig. 8A**) in the asymmetric unit that is very likely to be a physiologically relevant assembly of the protein based on several lines of evidence described below. All four subunits form a long, continuous α-helix with an equivalent conformation (**Fig. 8B**) that starts at residues 34-37 and ends at residues 87-91 in the different subunits. At the C-termini of these α-helices, the polypeptide chains could be traced into weak electron density through residues 98-101, but there is no interpretable electron density for the remaining 27 residues in any protomer. This entire segment of the protein has a high probability of backbone disorder according to the program DISOPRED3 (23), which predicts that over half of these disordered residues will participate in interprotein interactions. There is substantial amount of diffuse electron density in the crystal structure near the C-termini of the protomers that cannot be modeled in any specific conformation but that presumably derives from this disordered protein segment. The inability to model this density accounts for the relatively high R-factors of the refined structure (R_work_=30.8, R_free_ = 38.4). However, the other measures of refinement quality are all good (**Table S3**), and the close match between the refined backbone B-factors and the probability of backbone disorder according to Disopred3 (**Fig. 8C**) further supports the high quality of the refinement.

The core of the YhcB homotetramer is an antiparallel coiled-coil 4-helix bundle formed by residues 65-83 in each protomer (**Fig. 8A**). The interhelical packing pattern characteristic of coiled-coil interactions is interrupted by the alanine at position 73, which is responsible for the hole in the molecular surface visible in the view at the lower right in **Fig. 8A**, but the register of the coiled-coil interactions between the helices is nonetheless continuous through this region. This tetramer represents a dimer of V-shaped dimers that make parallel coiled-coil packing interactions at their N-termini spanning residues 37-51 (*i.e*., the closed end of the V). The subunits in this dimer splay apart starting at glutamine 54, which enables the open ends of the V-shaped dimer to interact to form the antiparallel coiled-coil 4-helix bundle. The overall assembly thus combines parallel and antiparallel coiled-coil packing interactions to form a tetramer with 222 symmetry (*i.e*., three orthogonal two-fold axes that intersect at the center of the assembly in the hole in the antiparallel coiled-coil region formed by alanine 72). While mixed parallel/antiparallel coiled-coil a-helical bundles have been observed before (*e.g*., in PDB id 4cq4 (24)), the program DALI (25) identifies the YhcB homotetramer as a novel protein structure because it has a unique tertiary structure in the region linking the parallel and antiparallel α-helical bundles.

The physiological relevance of this tetrameric assembly is supported by several lines of evidence, including strong evolutionary couplings (26) or pairwise evolutionary sequence correlations between the amino acids interacting in the central antiparallel coiled-coil bundle (**Fig. 8D**). The reliability of this computational analysis is supported by detection of the expected pattern of couplings between residues 3-4 apart in the long α-helix observed in the crystal structure. The strongest cluster of interactions in this analysis is between residues in the packing core of the antiparallel coiled-coil bundle, and couplings of this kind generally derive from direct physical contacts in a protein structure (27). While *E. coli* YhcB was not found to self-associate in our B2H screens nor in our co-purification experiments, Li *et al*. (2012) did find a self-interaction in a B2H screen using a different construct geometry. Detection of productive B2H interactions can depend on construct design due to the complexities of molecular geometry, especially for homo-oligomers, so at least some of the B2H data support physiologically significant self-interaction. Finally, the program PISA (28) also identifies the tetramer as a likely physiological oligomer based on quantitative analysis of its intersubunit packing interactions. Each subunit buries an average of 2,530 Å^2^ of solvent-accessible surface area in interfaces in the tetramer (755 Å^2^ in the parallel coiled-coil interface and 790 Å^2^ and 988 Å^2^ in the antiparallel coiled-coil interfaces), which is in the range characteristic of physiological oligomers.

While these observations all support the physiological significance of the tetramer observed in the crystal structure of *H. ducreyi* YhcB, the observation of a primarily monomeric structure in the crystallization stock suggests the affinity of the tetramer is such that it may reversibly dissociate *in vivo* dependent on local concentration. The absence or presence of binding partners that have higher affinity for the tetramer than the monomer could also modulate tetramer formation *in vivo*. The failure to detect self-association in our co-purification experiments is also consistent with relatively facile dissociation of the physiological tetramer.

Based on the location of its N-terminal transmembrane α-helices, the YhcB tetramer is likely to sit like an ~120 Å long handle parallel to the inner surface of the cytoplasmic membrane (lower left in **Fig. 8A**). The surface of this handle is characterized by a spiral pattern of strongly negative electrostatic potential (right in **Fig. 8A**) that is likely to influence YhcB’s interprotein interactions as well as its interactions with the nearby negatively charged surface of the cytoplasmic membrane. This structure could serve as a reversibly forming assembly point for multiprotein complexes on the surface of the membrane dependent on the local concentration of YhcB.

### The interaction sites of YhcB are conserved

We used site-directed mutagenesis to map and identify the residues involved in PPIs of YhcB. We divided the *E. coli* YhcB protein into 6 different regions based on the conserved residues identified by multiple sequence alignment and ConsurfDB analysis (**Fig. 9A**). The resulting variants cover different stretches of *yhcB* that we named v1 (N-terminal) to v6 (C-terminal). We also included a mutant lacking a transmembrane region (v7 or only cytoplasmic/CY) in order to investigate what role membrane localization (or the TM region) plays in the proper functioning of YhcB (v7 had the N-terminal 21 amino acids deleted).

**Figure 9.**
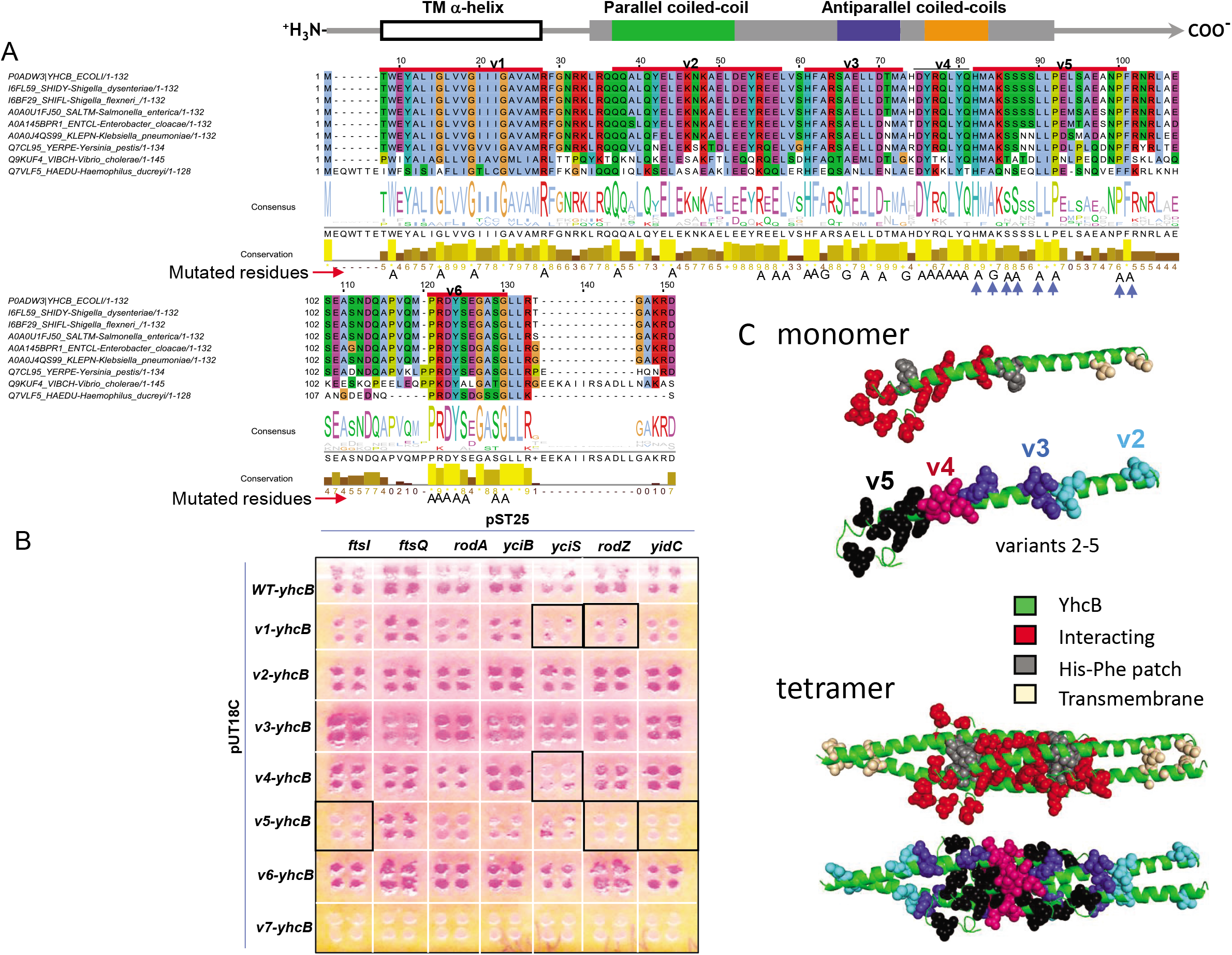
Interaction sites on YhcB. **(A)** Multiple sequence alignment of YhcB homologs across proteobacteria. The conserved residues are shown as a motif logo and histogram under the alignment, while a schematic of the 2° structure of *H. ducreyi* YhcB matching the depiction (**Fig. 8**) is shown above the alignment. The sequence is divided in to 6 different regions starting from v1 (N-terminus) to v6 (C-terminus), as indicated above the alignment. The highly conserved residues were mutated as shown beneath the sequence. **(B)** Bacterial two hybrid screens with YhcB mutants show the loss of specific interactions. The YhcB variant v5 showed the maximum loss in interactions with prey partners FtsI, RodZ and YidC. The v5 region possess several conserved residues predicted important for coiled-coil interactions as shown by arrow underneath the sequence in A. No interactions were detected in absence of the TM region (v7). **(C)** Protein models show mutated and thus potentially interacting residues in both YhcB monomer and tetramer.

Only the conserved residues of these regions were mutated (**Fig. 9A; mutated residues**). Each YhcB-variant had between four to eight amino acid substitutions and all residues were replaced with either alanine or glycine. In total, we created 37 mutations and each *yhcB* variant was tested against the positive interacting partners identified previously in B2H screens. The amino acids substitutions of *yhcB* variants v1, v4, and v5 had the strongest effect on interactions and were thus considered as potential PPI sites of YhcB (**Fig. 9B**). Amino acids H76, A78, S80, S81, L84, P86, P94, and F95 of YhcB-v5 (shown as arrowheads in **Fig. 9A**) seems to form an interaction site for multiple interacting proteins, especially FtsI, RodZ, YciS, and YidC (**Fig. 9B**). YhcB-v1 includes the conserved residues in the TM region only. These residues seem to be required for interactions with YciS and RodZ. The rationale for substitution of TM residues was to test if the region had any effect on PPIs or whether it was only required for interactions with the membrane. Interestingly, the TM region is required for interactions with all proteins: when it is deleted, all interactions are lost (v7 in **Fig. 9B,** but see Discussion). Notably, the substitutions in *yhcB*-v3 appear to result in several stronger interactions (**Fig. 9B**). The locations of these mutations are indicated in the monomer and tetrameric models we derived from the structure (**Fig. 9C**).

## Discussion

### Phenotypes and interactions

In *E.coli, yhcB* is conditionally essential and required for survival at high and low temperature, which is supported by previous large scale screens (29, 30). While the mechanisms underlying the temperature-related phenotypes remain unclear, heterologous expression of a *Caenorhabditis elegans* heat shock protein (CeHSP17) enabled *E. coli* cells to grow at 50°C and was cross-linked and co-purified with YhcB (31), linking YhcB to the *E. coli* heat shock response. Notably, we also observed an interaction between YhcB, YciS, and HemX proteins. YciS is a heat shock-induced protein (32) which has been co-purified with YhcB and HemX (33).

In *E. coli* and *Salmonella*, YhcB expression was reduced significantly upon overexpression of SdsR, a small RNA transcribed by the general stress sigma factor σS (34, 35). It was proposed that SdsR-mediated *yhcB* repression may be the primary cause for the SdsR-driven cell lysis because of the perturbation of cell division. These authors have reported defective growth with filamented cells upon *yhcB* deletion (13, 35) and support our results.

Sung et al. 2020 showed that yhcB deletions were restored by overexpressing YhcB protein, even when the transmembrane segment was missing. Effective complementation excludes the possibility that the phenotype was caused by polar effects of the deletion mutants or independent mutations elsewhere in the genome. While the phenotypes found by Sung et al. 2020 are similar to ours, most differences can likely be explained by somewhat different conditions and different strains (*E. coli* K-12 BW25113 in the Keio deletions used by us, but MG1655 used by Sung et al. 2020).

### Envelope stress-related interactions

YhcB physically interacts with outer membrane stress sensor proteases (*degQ* and *degS*) (**Table 3**) and both YhcB and DegS were predicted to be required for colonization of a host by *Vibrio* (36). Further, both DegQ and DegS proteases are involved in protein quality control in the cell envelope (37), suggesting a role of *yhcB* in stress related processes during cell-wall biogenesis or in cell envelope integrity. Also, in *E. coli*, the *yhcB* gene is predicted to be a part of MazF regulon and its mRNA is processed by MazF, a stress-induced endoribonuclease that is involved in post-transcriptional regulatory mechanism of protein synthesis globally in different stress-conditions (38).

The hypersensitivity of Δ*yhcB* to cell-wall acting antibiotics (14), specifically to vancomycin, could be because of impaired cell-wall biogenesis that leads to a permeable cell envelope (**Fig. S3-c**) and is further supported by the involvement of *yhcB* as part of the secondary resistome against colistin, an antibiotic targeting the outer membrane, in *Klebsiella pneumoniae* (39).

### Role in cell division and/or envelope biogenesis

A functional cell envelope and peptidoglycan biosynthesis is essential for cells to attach and form mature biofilms (40) and thus directly or indirectly support *yhcB* cell-wall associated phenotypes. The hypersensitivity of Δ*yhcB* cells towards cell-wall antibiotics in stationary phase potentially indicates an adaptive role during the stationary phase of bacterial cells. This notion is further supported by increased gene expression of YhcB during stationary phase growth in *Salmonella* (34) and in *E. coli* (41) and the inability of Δ*yhcB* to reduce length growth during stationary phase.

The PG-labelling using NADA and ED-DA fluorescent probes that report on PG synthesis show that lateral and septal PG synthesis is functioning globally as in wild type cells, apart from the positions of diffuse Z-ring localization. This suggests that YhcB is likely not directly involved in PG synthesis. However, a synthetic lethal and a physical interaction was observed between YhcB and YciB (**Fig. 7A-B**), a protein previously shown to be involved in PG synthesis (42) and a predicted intracellular septation protein (43). The deletion of *yhcB* does not only result in filamentation but also diffuse localization of Z-rings in those filamented cells. These phenotypes, together with the genetic and physical interactions of YhcB with FtsI, FtsQ, FtsZ, RodA, RodZ, and MreB, strongly support its role in cell division.

In order to accommodate our own and other observations, we propose a model for YhcB’s role in cell division which is based on previous models (44) (**Fig. 10**). YhcB interacts with several division proteins, including RodZ and RodA, suggesting that it is involved in the elongasome. Mid-cell localization of RodZ was shown to be essential for Z-ring formation (45). RodA forms a permanent complex with PBP2 (46) which was shown to be initially present at mid cell during Z-ring formation (47). The combined interactions of YhcB suggests that the elongasome brings YhcB to the assembly site of the divisome during preseptal PG synthesis. The divisome is a highly dynamic complex, hence its isolation has been only partly successful (47) but YhcB was detected as one of the protein of divisome complex isolated from cells in exponential and stationary phase using mass spectrometry (47).

**Figure 10.**
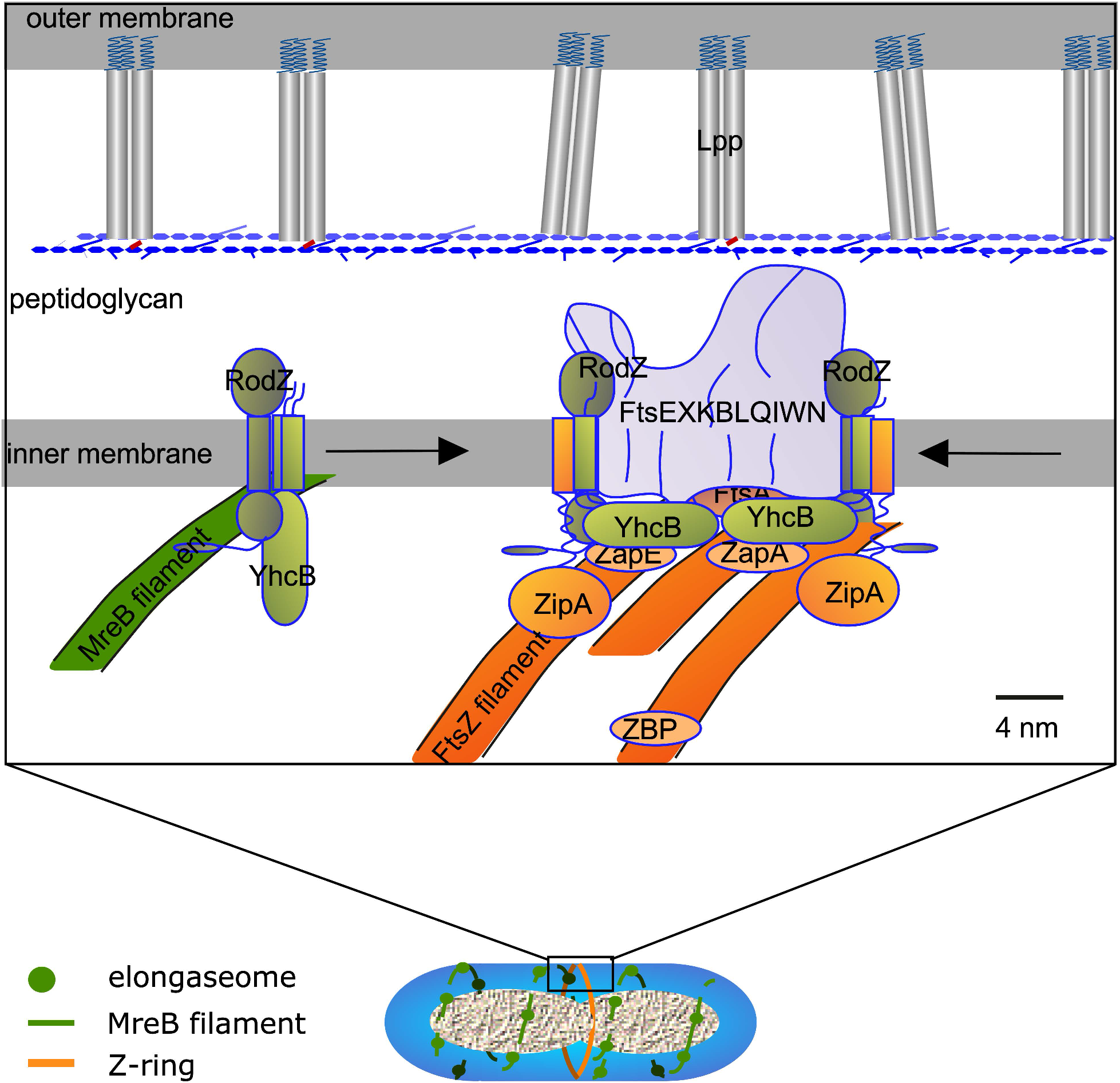
Model for YhcB function in cell division and Z-ring width maintenance. YhcB interacts as a dimer with RodZ that is part of the elongasome (sphere on green filament in cell schematic below). During peptidoglycan synthesis MreB (green filament) moves perpendicular to the length axes underneath the cytoplasmic membrane. Some of these filaments close to mid cell will be stalled by the Z-ring in the nascent state (orange). While some of the elongasome proteins will be involved in preseptal peptidoglycan synthesis on the periplasmic side of the cytoplasmic membrane, RodZ and YhcB interact with FtsZ filaments. As YhcB is likely present on both sides of the Z-ring the two dimers can associate into the tetrameric complex as observed by crystallography. This produces a bridge of ± 12 nm that can have multiple interactions with divisome proteins (here combined in one grey structure, “FtsEXKBLQIWN”) as observed by BTH. The Z-ring is formed by many filaments (with about 20 subunits each) that are connected by various FtsZ binding protein (ZBP, ZapA and ZapE) and linked to the cytoplasmic membrane by FtsA and ZipA (and possibly YhcB). With an average width of about 10 nm the Z-ring is of similar size as the RodZ-YhcB complex.

Consequently, many proteins have been reported that are supposed to help the FtsZ filaments to interact with each other (48, 49). But how are the boundaries of the Z-ring constrained? On the periplasmic side of the cytoplasmic membrane (CM), preseptal PG synthesis is thought to provide the borders in between which the new septum should be synthesised (50, 51). We suggest that YhcB helps to provide this function on the cytoplasmic side of the CM. While associated with RodZ at elongasome positions YhcB may be dimeric or monomeric but these interactions are dynamic and likely transient. When the elongasome is stalled at the nascent Z-ring from both sides of the ring, YhcB could come sufficiently close to form a weakly interacting tetramer parallel to the surface of the cytoplasm. This would provide a restricted width of the Z-ring of about 120 Å, which correlates well with the average width of the Z-ring of ±115 Å (52) and suggests that YhcB helps to determine the width of the Z-ring. The surface of the coiled coil of YhcB is sufficiently charged to interact with the membrane as well as with a number of cells division proteins and may tether the assembly in the close proximity of the CM.

### Structural considerations

The crystal structure of the *Haemophilus ducreyi* ortholog (**Fig. 8**) shows that its interaction sites cluster near the antiparallel alpha-helical coiled-coil at the center of the YhcB tetramer (**Fig. 9C**). Therefore, when the local concentration of YhcB is sufficient to drive homo-tetramerization, the tetramer and its 222 symmetry will enable it to mediate specific interactions tethered directly to the inner-surface of the cytoplasmic membrane. These interactions could serve as a focal point for organization of geometrically-defined supramolecular complexes controlling membrane morphology and dynamics during cell division. At lower concentrations, the monomer of YhcB could alternatively sequester the interaction interfaces of binding partners in a dissociated state in order to drive membrane morphology and dynamics in a different direction. The data presented in this paper supports YhcB playing a role in envelope biogenesis/integrity and cell division in Gamma proteobacteria. Biophysical studies of the interacting complexes identified in this paper, including cryo-EM reconstructions of the membrane-bound complexes, should provide deeper and more specific insight into the details of the related molecular mechanisms.

## Materials and Methods

### Bacterial strains and reagents

All strains used are listed below in their context of use. Strains were grown in LB media at 37°C unless otherwise mentioned. The Knock outs (KOs) were obtained from the *E. coli* Keio collection (53). PCR was used to confirm the *E. coli* Keio KOs using gene specific primers. *E. coli* TOP10 and DH5α were used for cloning. For protein expression, *E. coli* BL21(pLys) cells were used. *E. coli* was selected at 100 μg/ml ampicillin and/or 35 μg/ml chloramphenicol for expression in liquid media. All the expression experiments were done at 30°C unless otherwise mentioned. Antibiotics A22 and Mecillinam were purchased from Sigma-Aldrich (now Millipore Sigma).

### Phylogenetic analysis and Comparative genomic analysis

To determine potential for conservation of genes coding for our proteins of interest across bacterial species, we used the following methods. Starting with each gene’s UniProtKB identifier for *E. coli* K12, we identified membership of each in an orthologous group (OG) as defined by EggNOG v5.0 (54). Gene names, UniProtKB IDs, and corresponding EggNOG OGs are as follows: *ftsI* (P0AD68, COG0768), *ftsQ* (P06136, COG1589), *ftsZ* (P0A9A6, COG0206), *rodA* (P0ABG7, COG0772), *rodZ* (P27434, COG1426), *yciS* (P0ACV4, COG3771), *yhcB* (P0ADW3, COG3105), *yidC* (P25714, COG0706). In each case, the OG based on the broadest taxonomic definition was used (i.e., a COG). We then assembled a tree of 197 bacterial species and strains based on their NCBI taxonomy (55) and, for each, determined presence of at least one gene with membership in each of the above OGs as per EggNOG. Presence of these OG members was mapped and visualized with the iTOL tool v4 (56).

Genomic co-localization analysis was performed using the SEED annotation environment across representative members of sequenced bacterial species (57).

### Gateway cloning

Gateway cloning was performed according to instructions provided by the manufacturer (Invitrogen). The ORFs as entry clones for test proteins were obtained from the *E. coli* ORFeome clones assembled into the pDONR221 vector system (58). Then, the attL-flanked ORFs were cloned into the Gateway-compatible, attR-flanked bacterial two*-*hybrid (BACTH)-DEST plasmids (pST25-DEST, pUT18C-DEST, and pUTM18-DEST) using the LR reaction to generate attB-flanked ORFs in expression vectors. The plasmid preparations were done using Nucleospin column kits (Macherey Nagel). For the details of the B2H vectors and protocol, please refer to (59, 60).

### Bacterial Two Hybrid screening

Bacterial two hybrid screens were carried out as described in Mehla *et al*., 2017a. Briefly, the expression constructs of test proteins encoding the T25-X and T18-Y fusions were co-transformed into an adenylate cyclase (cya) deficient *E. coli* strain (BTH101). The competent cells were prepared using standard protocols (61). The co-transformants were selected on LB plates containing 100 μg/ml ampicillin and 100 μg/ml spectinomycin at 30°C after 48 hours. The selected co-transformants were screened on indicator plates at 30°C for 36-48 hours. The positive interactions were detected by specific phenotypes on indicator plates, i.e., blue colonies on LB-X-Gal-IPTG or red on MacConkey-Maltose medium. For quantification of PPIs (where required), the β-galactosidase assay was used (62). The details of test proteins are shown in **Table S4**.

### Affinity purification combined with mass spectrometry and genetic crosses

YhcB fused to SPA-tag, chromosomally at the C-terminus, was confirmed by immunoblotting using anti-FLAG antibody, and then purified in the presence and absence of various mild non-ionic detergents, essentially as described (18). The stably-associated proteins were detected by MS using the SEQUEST/ STATQUEST algorithm, following established procedures (18, 33). Genetic crosses were conducted as previously described (33) by conjugating Hfr Cavalli (Hfr C) *yhCB::Cm^R^* donor gene deletion mutant marked with chloramphenicol against the select set of F-‘recipient’ non-essential single gene deletion or essential hypomorphic mutants marked with kanamycin resistance, including functionally unrelated gene *JW5028* (63) from the Keio single gene deletion mutant library, to generate digenic mutants after both antibiotic selection.

### Mapping Protein-protein interaction site (s): Mutagenesis of *yhcB*

To map interaction site(s), mutants of YhcB were constructed. YhcB was divided into 6 different regions and in each region 3-4 site-specific substitutions were inserted. Also, a cytoplasmic version without the TM region of YhcB was constructed. Only conserved residues of YhcB were mutated (as shown in **Fig. 9A**). Mutant DNA sequences encoding specific mutants were synthesized as full gene sequences by Geneart (ThermoFisher pvt Ltd). These sequences were further cloned into pDNOR/Zeo using the BP Clonase reaction of Gateway cloning (Invitrogen). The transformants with correct sequences were confirmed by sequencing at least 2 different clones. The ORFs were further sub-cloned into bacterial two hybrid vector pUT18C followed by co-transformation and screening for interactions against prey proteins as discussed above (**Section B2H**).

### Growth Inhibition/sensitivity against drugs

The growth of both WT and Δ*yhcB* strains was monitored in different media and in different conditions, such as different carbon sources, antibiotics as well as rich and selective media, each in 96-well microplates at 37 °C. The bacterial growth was measured as the optical density (OD) at 562 nm using a plate reader. The % inhibition (or survival) was calculated as previously described (64).

### Antibiotic susceptibility testing (Serial dilution assay)

An overnight culture of *E. coli* strains (both WT and Δ*yhcB*) was tested for susceptibility towards cell-wall antibiotics using serial dilutions. 10^7^ cells/ml were serially diluted, and 5 μl of each dilution was spotted on LB with or without added antibiotic or other compounds (e.g 1% carbon sources). For MacConkey plates, 3 μl of each dilution was used. The plates were then imaged after 24 hours or at other specific time points (see text for details). A22 (1 μg/ml) or Mecillinam (0.12-0.25 μg/ml) was used in dilution assays on hard agar media. These concentrations were chosen based on effective ranges tested by Nichols et al. 2011 (0.5, 2, 5, and 15 μg/ml for A22, resulting in [log] reductions of growth by −1.015628, −4.344713, −3.311473, −3.978085), and Mecillinam (0.03, 0.06, 0.09, and 0.12 μg/ml, resulting in [log] reductions of −0.339263, −4.244134, −8.923793, −6.08356, respectively).

### Persister/survivor cell assay

Persister/survivor cell assays were done as reported previously (65). Persistence was determined by determining the number of colony-forming units (CFUs) upon exposure to A22 (1 μg/ml) and Mecillinam (0.12 μg/ml). We determined the number of persister/survivor cells in the Δ*yhcB* strain upon exposure to cell-wall antibiotics for 6 hours. The overnight culture was sub-cultured at 37°C for 2 hours and the cells in early log phase were treated with antibiotics. The overnight cells were used as stationary phase cells. For determination of CFUs, 2 μl of culture (10^7^ cells/ml) was resuspended in fresh medium, serially diluted, and plated on solid LB medium. The number of survivor cells were determined as colony forming units (CFUs) upon antibiotic treatment. The CFUs were expressed as % survival of treated vs untreated cells.

### FtsZ localization

The FtsZ ring formation and localization was monitored using both immunolabelling and GFP fusion of FtsZ. The Δ*yhcB* and its parental strain BW25113 (WT) were grown in LB at 37 °C for 24 h (ON), then dilute 1:1000 and grown to an OD_650_ nm of 0.3 (EXP) or to and OD_650_ nm of 1.2 (STAT), fixed for 15 min by addition of a mixture of formaldehyde (f. c. 2.8%) and glutaraldehyde (f. c. 0.04%) to the cultures in the shaking water bath and immunolabeled as described previously (66) with Rabbit polyclonal antibodies against FtsZ (67). As secondary antibody, donkey anti-rabbit conjugated to Cy3 or to Alexa488 (Jackson Immunochemistry, USA) diluted 1:300 in blocking buffer (0.5% (wt/vol) blocking reagents (Boehringer, Mannheim, Germany) in PBS) was used, and the samples were incubated for 30 minutes at 37°C. For immunolocalization, cells were immobilized on 1% agarose in water slabs coated object glasses as described (67) and photographed with an Orca Flash 4.0 (Hamamatsu, Japan) CCD camera mounted on an Olympus BX-60 (Japan) fluorescence microscope through a 100x/*N.A*. 1.35 oil objective. Images were taken using the program ImageJ with MicroManager (https://www.micro-manager.org). Phase contrast and fluorescence images were combined into hyperstacks using ImageJ (http://imagej.nih.gov/ij/) and these were linked to the project file of Coli-Inspector running in combination with the plugin ObjectJ (https://sils.fnwi.uva.nl/bcb/objectj/). The images were scaled to 15.28 pixel per μm. The fluorescence background has been subtracted using the modal values from the fluorescence images before analysis. Slight misalignment of fluorescence with respect to the cell contours as found in phase contrast was corrected using Fast-Fourier techniques as described (67). Length, diameter and fluorescence concentration were measured using Coli-Inspectror running in combination with the plugin ObjectJ (https://sils.fnwi.uva.nl/bcb/objecti/) as described (67).

For GFP tagged FtsZ localization, the cells were grown at 37°C in LB media to exponential phase. Imaging was done on M16 glucose plus casamino acids pads with 1% agarose at room temperature. Phase contrast images where collected on a Nikon Eclipse Ni-E epifluorescent microscope equipped with a 100X/1.45 NA objective (Nikon), Zyla 4.2 plus camera, NIS Elements software (Nikon). A functional FtsZ fusion was made by inserting msfGFP at an internal site of FtsZ and replacing the native copy of FtsZ with the fusion protein.

### Peptidoglycan (PG)-labelling and localization

The PG labelling studies were conducted as previously reported (15, 16). Briefly, overnight cultures were started from single colonies grown from −80°C freezer stocks (plated overnight). Experimental cultures were then started in 5 ml of LB. Double the amount of the wild type strain was used to inoculate cultures for the *yhcB* mutant (50 μl vs 100 μl in 5 ml) in order to attain ODs as close as possible after two and a half hours of growth (OD_600_ values of 0.8 and 0.7, respectively). This was done to minimize the time required to back-dilute and achieve exactly equivalent OD readings, which likely would have had an effect on the rate of PG synthesis / and turnover.

We took logarithmic growing cultures (WT in LB and Δ*yhcB* in LB + 1% glucose) and conducted a short pulse with our 1st gen probes (NADA) 2^nd^ gen probes (EDA-DA) for 45 seconds. Glucose supplementation was utilized in the Δ*yhcB* culture in order to ensure each strain achieved comparable growth kinetics. After the short pulse, bacteria cultures were fixed immediately in 70% (final concentration) ice-cold ethanol for 20 minutes. NADA-labeled cells were washed three times in PBS, mounted on 1% agar pads, and imaged via a Zeiss 710 confocal laser scanning microscope. EDA-DA-labeled cells were subsequently bound to azide-conjugated Alexa Fluor 488 via a click chemistry reaction using a Click-iT Cell Reaction Buffer Kit (Invitrogen), as previously described (16). Cells were then washed three times in PBS +3% BSA, once in PBS, mounted on 1% agar pads, and imaged via Zeiss Elyra PS1 super resolution microscope in structured illumination (SIM) mode. Images are representative of 20 fields of view observed per condition / strain examined.

### Light microscopy and image analysis

The cells were stained and imaged to visualize cell membrane and nucleoid using FM4-64 SynaptoRed™ C2 (FM4-64 (4-[6-[4-(Diethylamino) phenyl]-1,3,5-hexatrien-1-yl]-1-[3-(triethylammonio) propyl] pyridinium dibromide, Biotium Inc.) and DAPI, respectively. The cells were imaged on an Olympus BX41 microscope at 100x in a dark room. Images were captured with a microscope digital camera (AmScope MU1400). ImageJ software was used for measuring cells dimensions/length (68).

### Protein expression, purification, and light-scattering analysis

Residues 31-128 from the YhcB ortholog in *H. ducreyi* (HD1495, UniProt id Q7VLF5, Northeast Structural Genomics Consortium target HdR25) were cloned into a pET21-derived T7 expression vector between an N-terminal initiator methionine residue and a C-terminal affinity tag with sequence LEHHHHHH, and this vector was deposited at the ASU Biodesign Institute (http://dnasu.org/DNASU/GetCloneDetail.do?cloneid=338479). Cloning, purification, and quality-control analysis methods were described previously (69). In brief, after growing cells to logarithmic phase at 37 °C in chemically defined MJ9 medium with 0.4% (w/v) glucose, protein expression was induced overnight at 18 °C with 1 mM IPTG. Soluble protein was purified by Ni-NTA chromatography followed by Superdex 75 gel-filtration in 100 mM NaCl, 5 mM DTT, 20 mM Tris•HCL, pH 7.5. Pooled fractions were ultrafiltered in an Amicon device prior to flash-freezing in liquid N2 in single-use aliquots at crystallization concentration. Protein quality was characterized using SDS-PAGE, MALDI-TOF mass spectrometry (12,574.8 daltons observed *vs*. 12,549.6 predicted for selenomethionine-labeled wild-type protein), and size-exclusion-chromatography/multiangle-light-scattering (SEC-MALS) in the gel filtration buffer using a Shodex KW802.5 column (Showa Denko, New York, NY) with a Wyatt Technology (Santa Barbara, CA) detector system (**Fig. S5**).

### Protein crystallization, x-ray structure determination, and refinement

Crystallization screening and optimization were performed using the microbatch method under paraffin oil (70, 71). The structure was solved using single-wavelength anomalous diffraction phasing (72) of a selenomethione-labeled construct harboring I51M and L72M mutations, which crystallized similarly to the wild-type construct. These mutations were introduced to increase selenomethionine phasing power compared to the WT construct, which only has a single N-terminal methionine that is disordered in the crystal structure. The mutations were introduced at uniformly hydrophobic positions that show methionine in some orthologs in an YhcB sequence alignment, based on the premise that such positions are likely to be at least partially buried and therefore well-ordered and provide good phasing power. Diffraction data were collected at 100 °K on beamline 19-ID at the Advanced Photon Source using x-rays at the Se K-edge (λ = 0.979 Å) and processed using HKL2000 (73). The structure was solved and refined at 2.8 Å resolution using PHENIX (74), built using interactive cycles in Coot (75), validated using PROCHECK (76), and deposited in the RCSB Protein Data Bank under accession code 6UN9. Data collection and refinement statistics are shown in **Table S3**.

The relatively high free R-factor for a structure at this resolution (38.4%) is attributable to the low mean intensity of the diffraction dataset (<I/σ_I_> = 4.1>) combined with the high degree of disorder in the crystallized construct (**Table S3**). Over 30% of residues are disordered and could not be modeled at all, while greater than 10% of the residues are only partially ordered, preventing accurate modeling with a single coordinate model with individual atomic B-factors. The disordered residues and the refined B-factors of the modeled residues (**Fig. 3C**) both correlate very closely with the probability of backbone disorder calculated by the program DISOPRED3 (23), which uses exclusively primary sequence data and is therefore completely independent of the crystal structure. Furthermore, the accuracy of the structure solution and refined coordinate model are supported by four additional factors, all of which are independent of one another and the backbone disorder prediction. First, the interprotomer contacts in the structure correlate strongly with pairwise evolutionary couplings in the YchB protein family (**Fig. 3D**) as calculated by the program GREMLIN (80), which also uses exclusively primary sequence data and is completely independent the crystal structure. Second, an anomalous difference Fourier map calculated with the refined phases shows strong peaks at the positions of the selenium atoms in the engineered selenomethione residues in the protein construct and no significant peaks anywhere else in the unit cell (**Fig. S6A**). Third, the 2f_0_-f_c_ electron density map calculated from the refined coordinate model shows excellent agreement with the model consistent with the 2.8 Å overall resolution of the crystal structure (**Fig. S6B**). Finally, the crystallographically related tetramers fill the unit cell and make appropriate packing interactions to stabilize the modeled structure in the lattice, which has a 65% solvent content (**Fig. S6**).

### Protein structure analysis

Coiled-coil sequence propensity was analyzed using the program Coils (77), which indicates high probability of coiled-coil formation for residues 44-64, 37-75, and 30-82 for windows of 14, 21, and 28 residues, respectively. Coiled-coil packing interactions in the crystal structure were analyzed using Socket (78) and Twister (79). Buried solvent-accessible surface area was calculated using PISA (28). Backbone disorder probability was calculated using DISOPRED3 (23), and evolutionary couplings were calculated using Gremlin (80). Molecular graphics images were generated using PyMOL (https://pymol.org/2/), which was also used to calculate *in vacuo* surface electrostatics.

## Acknowledgements

We are thankful to Dr. Scot Ouellette (University of Nebraska) and Dr. Catherine Paradis-Bleau (Université de Montréal, Montreal, Quebec) for providing us the bacterial two-hybrid vectors and the pCB112 plasmid for β-galactosidase assays. Dr. Michael VanNieuwenhze (Indiana University) provided peptidoglycan labeling reagents (NADA and EDA-DA). This work was supported by the Natural Sciences and Engineering Research Council of Canada, Grant DG-20234 to M. B., National Institutes of Health grants 5U54GM094597 to G.T.M., and GM109895 to P.U. and a faculty start up award to G.L. The views expressed here are those of the authors and should not be construed as official or representing the views of the Department of Defense or the Uniformed Services University.

## Author contributions

JM carried out the B2H interactions, site-directed mutagenesis, and phenotypic, studies. GL and MB conducted the PG labelling and labelling study and wrote the corresponding section of the manuscript, RM and TdB did the FtsZ localization studies, JHC and JM did the phylogenetic analysis, AH, AG, SYK, SP and MB analyzed genetic and MS interactions data, NS helped with B2H screens, SV, RX, GTM, and JFH designed and purified YhcB protein constructs and solved the x-ray crystal structure, JM, JHC, JFH, and PU wrote the manuscript; GL, RM, MB and TdB edited the manuscript. PU analyzed data, secured funding, and wrote part of the manuscript.

## Supplementary Figures

**Fig. S1**. Gene synteny of *yhcB* and neighboring genes in selected proteobacterial genomes. Tree from iTol (56).

**Fig. S2.** Imaging of Δ*yhcB* cells. Δ*yhcB* is longer and thinner than its parental strain BW25113, but nucleoid topography seem to be normal. (**A**). Phase contrast images of the cells and DAPI fluorescence images of the nucleoids of BW25113 WT cells and Δ*yhcB* cells grown in minimal glucose medium (GB4) at 28°C. The scale bar equals 5 μm. (**B)**. Length and diameter of both strains grown in rich medium (LB with 5 g NaCl/L) at 37°C and Gb4 28°C. **(C)** Demographs of DAPI stained nucleoid distribution in BW25113 (n = 750) (first panel) and Δ*yhcB* (n = 650) cells grown in TY at 37°C (second panel), BW25113 (n = 1521, third panel) and Δ*yhcB* (n = 1095, fourth panel) cells grown Gb4 at 28°C, respectively. The cells are sorted according to cell length and the white outline based on the phase contrast images represents the length of the cells.

**Fig. S3.** Phenotypes associated with *yhcB* deletion.

**(a)** Growth curve/profile of Δ*yhcB* strain in LB and on LB agar. *yhcB* is required for optimal growth of *E. coli*. The Δ*yhcB* strain never reached an OD_562_ comparable to WT strain both in LB and LB-glucose. Data represents at least three independent experiments. (**b)** Serial dilution of Δ*yhcB* in LB and LB-glucose on hard agar plates shows similar growth patterns after 24 hours. (**c)** β-galactosidase (CPRG) assay. Cell envelope integrity of Δ*yhcB* strain was tested using a β-galactosidase assay. Both deletion of *yhcB* and an interactor Δ*yciS* showed pinkish colored cells showing defective or permeable cell envelope. **(d)** The Δ*yhcB* cells were found deficient in biofilm formation in LB media. The relative or fold difference in biofilm formation in Δ*yhcB* cells *vs* WT cells is shown here.

**Fig. S4.** Hypersensitivity of Δ*yhcB* strain

**(a).** The susceptibility of an Δ*yhcB* strain to antibiotics targeting the cell wall biogenesis reported in a phenomic profiling of *E. coli* screen.

**(b).** Hypersensitivity of 2 days old stationary phase Δ*yhcB* cells against cell-wall acting antibiotics. The Δ*yhcB* cells were found hypersensitive to A22 and Mecillinam. The 2-day old stationary phase cells were not able to grow in presence of A22 and Mecillinam. Top=A22, Bottom=Mecillinam

**Fig. S5.** Size exclusion chromatography and multi-angle light-scattering (SEC-MALS) analysis of the cytosolic segment from *H. ducreyi* YhcB. The analysis was performed on a Shodex KW802.5 column equilibrated in 100 mM NaCl, 5 mM DTT, 20 mM Tris•Cl, pH 7.5. The dotted horizontal lines indicate the predicted molecular weights for a monomer (12,550 daltons), tetramer, and hexadecamer. Quantitative analyses including integration of the refractive index trace (blue) indicate a total recovery of 386 μg of protein distributed between species with average molecular weights of 15.5 kDa (96.3%), 50.2 kDa (2.5%), 199 kDa (0.9%), and 3,520 kDa (0.2%). The molecular weight of the smallest species is 24% higher than the predicted molecular weight for a monomer of this protein construct (12.5 kDa), which could reflect a reversible tendency to oligomerize or, alternative, inaccuracy in the light-scattering-based molecular weight determination in this size range. Reversible oligomerization is concentration-dependent, which generally produces a characteristic parabolic trend in the estimated molecular weight across a SEC peak, with larger values at the center of the peak where the protein concentration is higher compared to is tails. Therefore, the consistency of the calculated molecular weight across the major elution peak suggests the discrepancy in measured vs. predicted molecular weight is more likely to be attributable to inaccurate calibration in this molecular weight range rather than reversible oligomerization. This analysis was performed on the selenomethionine-labeled wild-type protein construct comprising residues 31-128 with an N-terminal methionine and C-terminal affinity tag with sequence LEHHHHHH but without the L51M or L72M mutations used for selenomethionine phasing of the crystal structure.

**Fig. S6.** Electron density maps and lattice packing in the x-ray crystal structure of Haemophilus ducreyi YhcB. **(A)** Anomalous difference Fourier map calculated using phases from the final refined model of seleonmethionine-labeled *H. ducreyi* I51M-L72M-YhcB. The map contoured at 5 σ is shown in red, the refined atomic model of the tetramer in the asymmetric unit of the crystal structure is shown in blue line representation, its symmetry mates in the crystal lattice are shown in pale green line representation, and the boundaries of the unit cell are shown as yellow lines. The strong peaks in the anomalous difference Fourier map all correspond to selenium atoms in the side chains of the engineered residues Met-51 and Met-72. The latter residue adopts multiple conformations in some subunits in the physiological tetramer. **(B)** The same image in panel **A** but with the addition of the 2f_0_-f_C_ electron density map calculated from the final refined coordinate model shown in light blue contoured at 1.5 σ.

**Supplementary Tables**

**Table S1**.

YhcB PPIs detected by B2H and mass-spectrometry based proteomic screens.

**Table S2.**

**Cross-species protein-protein interactions of YhcB.** The cross-species interactions of YhcB were measured in proteobacteria *E. coli, Yersinia pestis* and *Vibrio cholerae*. We found several inter and intra-species conserved interactions of YhcB.

**Table S3.**

Crystallographic data from *Haemophilus ducrey* YhcB^1^.

**Table S4.**

Information, including sequence coverage and identity between homologs of the *E. coli, Yersinia pestis* and *Vibrio cholerae* proteins tested in B2H screens.

