## Supplementary Figures for "YhcB (DUF1043), a novel cell division protein conserved across gamma-proteobacteria"

**YhcB is conserved in proteobacteria**

YhcB is conserved across most gamma-proteobacteria but hardly present in any other bacterial clades (**Fig. 1**). Even within gamma-proteobacteria it appears to have been lost secondarily in many clades. While not highly conserved, its amino acid sequence in *E. coli* is 45% and 80% identical to its homologs in *Vibrio* and *Yersinia,* respectively.

In *Escherichia coli*, the *yhcB* gene is located upstream of two periplasmic outer membrane stress sensor (serine) proteases (*degQ* and *degS*) and predicted to be a member of the same transcriptional unit (**Fig. S1**). Upstream of *yhcB,* cell division gene *zapE* (*yhcM*) is encoded on the opposite strand, an arrangement that is conserved across many proteobacteria. Genomically, *yhcB* is clustered with genes involved in outer membrane stress-response and cell division pathways.

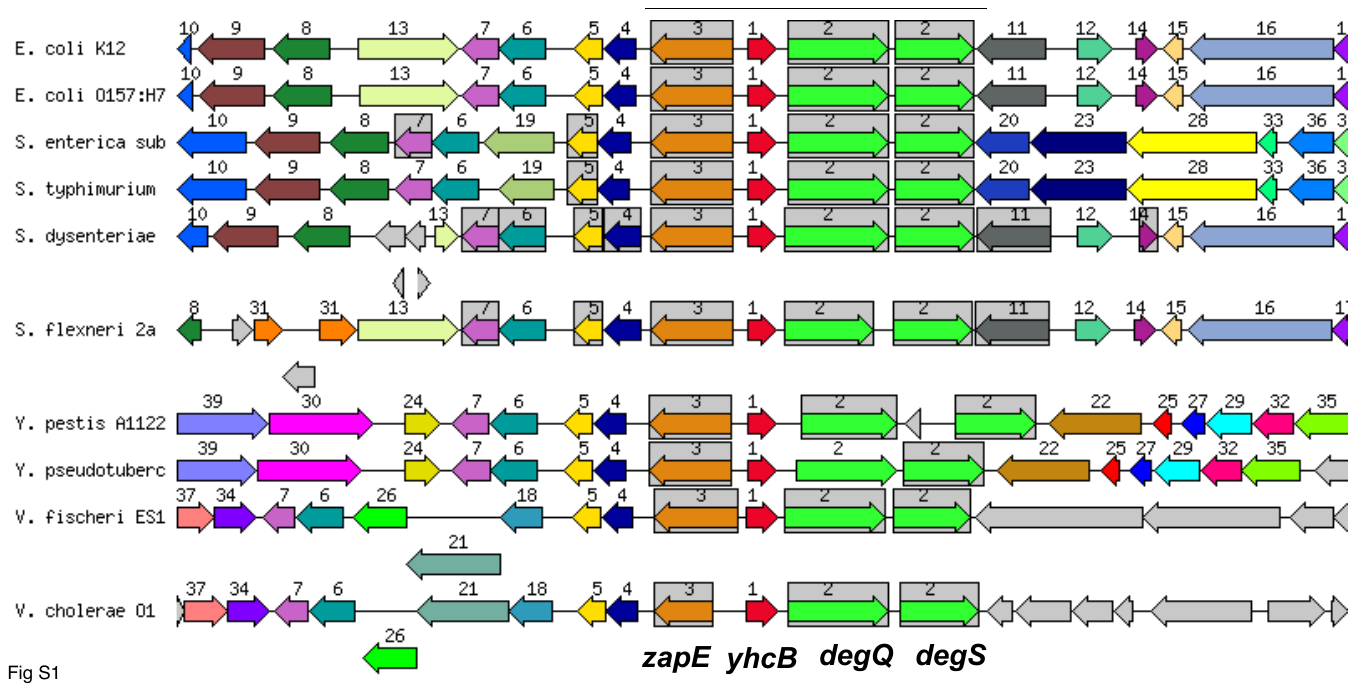

**Fig. S1.** Gene synteny of yhcB and neighboring genes in selected proteobacterial genomes. Tree from iTol (56).

**Phenotypes associated with *yhcB* gene deletion**

In order to understand the function and phenotypes of *yhcB*, we utilized a Δ*yhcB* deletion strain to carry out extensive phenotypic testing. In LB at 37°C, Δ*yhcB* grows with a mass doubling time of 25 min, whereas the WT grows with 22 min. Close to stationary phase the Δ*yhcB* cells starts to die. With an optical density difference of 0.3 OD units the difference in survival is more than 17-fold. In LB with 0.5 g salt per liter at 37°C and in minimal glucose medium at 28°C the Δ*yhcB* strain is slightly longer and thinner than the wild type parental strain (**Fig. S2**).

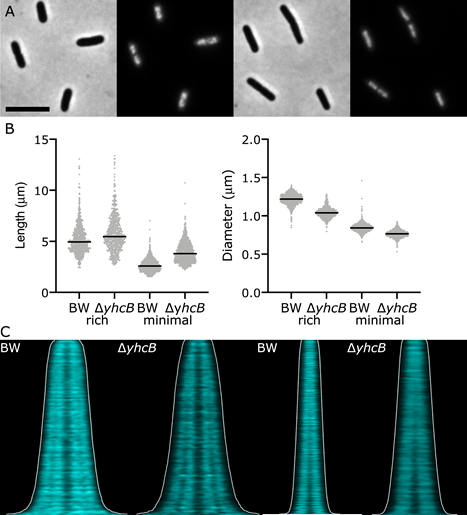

**Fig. S2.** Imaging of ∆*yhcB* cells. ∆*yhcB* is longer and thinner than its parental strain BW25113, but nucleoid topography seem to be normal. (**A**). Phase contrast images of the cells and DAPI fluorescence images of the nucleoids of BW25113 WT cells and ∆*yhcB* cells grown in minimal glucose medium (GB4) at 28°C. The scale bar equals 5 µm. (**B)**. Length and diameter of both strains grown in rich medium (LB with 5 g NaCl/L) at 37°C and Gb4 28°C. (C) Demographs of DAPI stained nucleoid distribution in BW25113 (n = 750) (first panel) and ∆*yhcB* (n = 650) cells grown in TY at 37°C (second panel), BW25113 (n = 1521, third panel) and ∆*yhcB* (n = 1095, fourth panel) cells grown Gb4 at 28°C, respectively. The cells are sorted according to cell length and the white outline based on the phase contrast images represents the length of the cells.

The Δ*yhcB* strain grows slowly compared to WT at 37°C and never reaches a max OD_562_ comparable to the parent strain **(Fig S3-a &b)**. Further, the cell envelope integrity in Δ*yhcB* cells was tested using a β-galactosidase assay as described previously (14). Briefly, the cells were transformed with a pCB112 plasmid by mating with an overnight culture of JA200/pCB112 donor strain. The Lac+ exconjugants were selected on LB-Kan-Cam plates and were tested on LB agar plates containing 20 µg/ml CPRG. A light pink color was observed in Δ*yhcB* cells indicating a break down in barrier function of the cell envelope (**Fig S3-c**). The Δ*yhcB* strain also exhibited significantly reduced biofilm formation (**Fig S3-d**). The biofilm formation was determined as described previously (https://www.jove.com/t/2437/microtiter-dish-biofilm-formation-assay). Briefly, the cells were grown in a 96-well plate and the biofilm formation was measured by staining with 0.1 % solution of crystal violet for 15 min at room temperature, followed by washing and quantification of biofilm. The biofilm was quantified by measuring the absorbance at 550 nm and normalized with respect to WT strain.

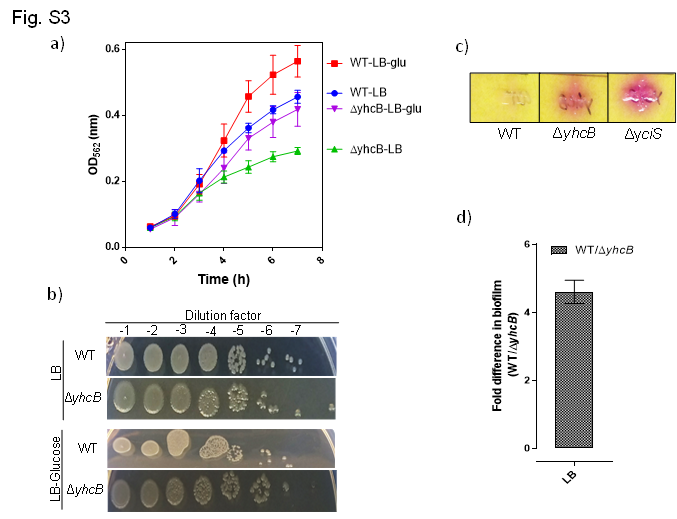

**Fig. S3.** Phenotypes associated with *yhcB* deletion.

**(a)** Growth curve/profile of ∆*yhcB* strain in LB and on LB agar. *yhcB* is required for optimal growth of *E. coli*. The ∆*yhcB* strain never reached an OD_562_ comparable to WT strain both in LB and LB-glucose. Data represents at least three independent experiments. (**b)** Serial dilution of Δ*yhcB* in LB and LB-glucose on hard agar plates shows similar growth patterns after 24 hours. (**c)** β-galactosidase (CPRG) assay. Cell envelope integrity of Δy*hcB* strain was tested using a β-galactosidase assay. Both deletion of *yhcB* and an interactor Δ*yciS* showed pinkish colored cells showing defective or permeable cell envelope. **(d)** The ∆*yhcB* cells were found deficient in biofilm formation in LB media. The relative or fold difference in biofilm formation in ∆*yhcB* cells vs WT cells is shown here.

Furthermore, Δ*yhcB* strain showed sensitivity to several cell-division/envelope biogenesis targeting/related antibiotics. Given that we found cell shape and cell division-related phenotypes (e.g. filamented) and susceptibility of Δ*yhcB* strain to several peptidoglycan (PG)-targeting antibiotics (**Fig S4-a**), we tested the effect of cell-wall targeting antibiotics (A22, and Mecillinam) on Δ*yhcB* cells. Proteins of the cell elongasome such as MreB and PBP2 are direct targets of the cell-wall antibiotics A22 and Mecillinam, respectively. A22 (S-[3,4-Dichlorobenzyl] isothiourea) destabilizes MreB filaments (20) and Mecillinam is a β-lactam that specifically inhibits PBP2 (21). We confirmed that the Δ*yhcB* strain was hypersensitive to both A22 and Mecillinam (**Fig 3D**). Furthermore, the ∆*yhcB* cells didn’t revive in presence of A22 and Mecillinam whereas ∆*yhcB* cells in absence of antibiotics behaved normally.

**Fig. S4.** Hypersensitivity of Δ*yhcB* strain

**(a).** The susceptibility of an Δ*yhcB* strain to antibiotics targeting the cell wall biogenesis reported in a phenomic profiling of *E. coli* screen.

**(b).** Hypersensitivity of 2 days old stationary phase ∆*yhcB* cells against cell-wall acting antibiotics. The ∆*yhcB* cells were found hypersensitive to A22 and Mecillinam. The 2-day old stationary phase cells were not able to grow in presence of A22 and Mecillinam. Top=A22, Bottom=Mecillinam

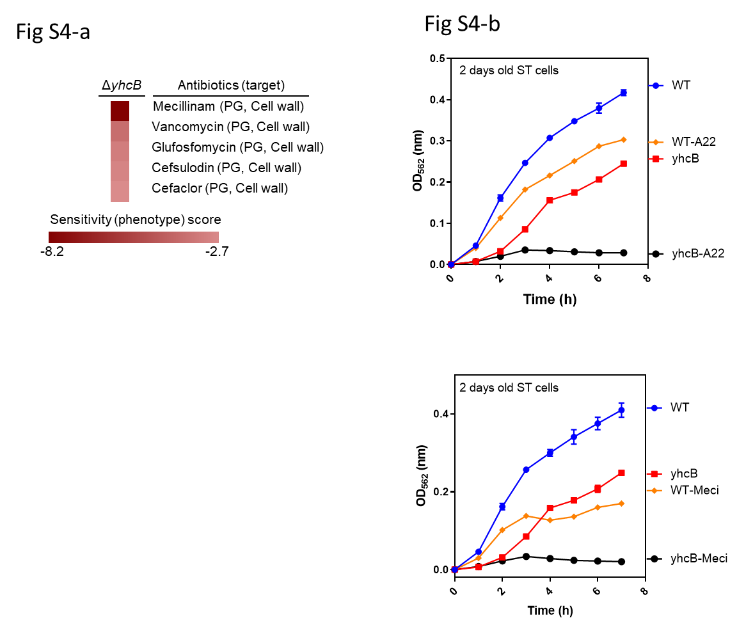

**Fig. S5.** Size exclusion chromatography and multi-angle light-scattering (SEC-MALS) analysis of the cytosolic segment from *H. ducreyi* YhcB. The analysis was performed on a Shodex KW802.5 column equilibrated in 100 mM NaCl, 5 mM DTT, 20 mM Tris•Cl, pH 7.5. The dotted horizontal lines indicate the predicted molecular weights for a monomer (12,550 daltons), tetramer, and hexadecamer. Quantitative analyses including integration of the refractive index trace (blue) indicate a total recovery of 386 µg of protein distributed between species with average molecular weights of 15.5 kDa (96.3%), 50.2 kDa (2.5%), 199 kDa (0.9%), and 3,520 kDa (0.2%). The molecular weight of the smallest species is 24% higher than the predicted molecular weight for a monomer of this protein construct (12.5 kDa), which could reflect a reversible tendency to oligomerize or, alternative, inaccuracy in the light-scattering-based molecular weight determination in this size range. Reversible oligomerization is concentration-dependent, which generally produces a characteristic parabolic trend in the estimated molecular weight across a SEC peak, with larger values at the center of the peak where the protein concentration is higher compared to is tails. Therefore, the consistency of the calculated molecular weight across the major elution peak suggests the discrepancy in measured vs. predicted molecular weight is more likely to be attributable to inaccurate calibration in this molecular weight range rather than reversible oligomerization. This analysis was performed on the selenomethionine-labeled wild-type protein construct comprising residues 31-128 with an N-terminal methionine and C-terminal affinity tag with sequence LEHHHHHH but without the L51M or L72M mutations used for selenomethionine phasing of the crystal structure.

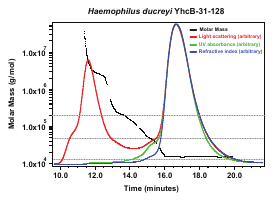

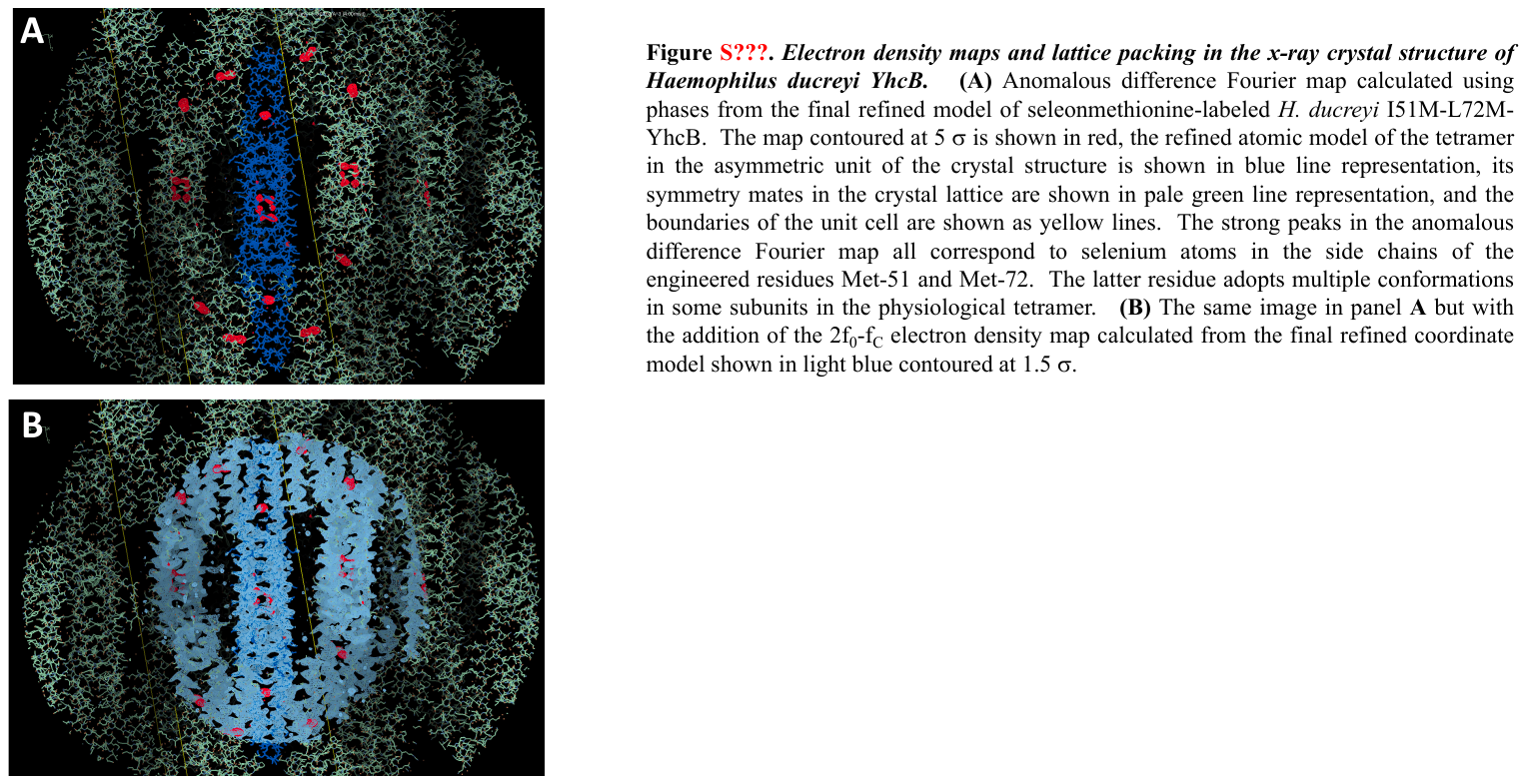

**Fig. S6.** Electron density maps and lattice packing in the x-ray crystal structure of Haemophilus ducreyi YhcB. **(A)** Anomalous difference Fourier map calculated using phases from the final refined model of seleonmethionine-labeled *H. ducreyi* I51M-L72M-YhcB. The map contoured at 5 σ is shown in red, the refined atomic model of the tetramer in the asymmetric unit of the crystal structure is shown in blue line representation, its symmetry mates in the crystal lattice are shown in pale green line representation, and the boundaries of the unit cell are shown as yellow lines. The strong peaks in the anomalous difference Fourier map all correspond to selenium atoms in the side chains of the engineered residues Met-51 and Met-72. The latter residue adopts multiple conformations in some subunits in the physiological tetramer. **(B)** The same image in panel **A** but with the addition of the 2f_0_-f_C_ electron density map calculated from the final refined coordinate model shown in light blue contoured at 1.5 σ.
